# Comprehensive analyses of a large human gut Bacteroidales culture collection reveal species and strain level diversity and evolution

**DOI:** 10.1101/2024.03.08.584156

**Authors:** Zhenrun J. Zhang, Cody G. Cole, Michael J. Coyne, Huaiying Lin, Nicholas Dylla, Rita C. Smith, Emily Waligurski, Ramanujam Ramaswamy, Che Woodson, Victoria Burgo, Jessica C. Little, David Moran, Amber Rose, Mary McMillin, Emma McSpadden, Anitha Sundararajan, Ashley M. Sidebottom, Eric G. Pamer, Laurie E. Comstock

## Abstract

Species of the Bacteroidales order are among the most abundant and stable bacterial members of the human gut microbiome with diverse impacts on human health. While Bacteroidales strains and species are genomically and functionally diverse, order-wide comparative analyses are lacking. We cultured and sequenced the genomes of 408 Bacteroidales isolates from healthy human donors representing nine genera and 35 species and performed comparative genomic, gene-specific, mobile gene, and metabolomic analyses. Families, genera, and species could be grouped based on many distinctive features. However, we also show extensive DNA transfer between diverse families, allowing for shared traits and strain evolution. Inter- and intra-specific diversity is also apparent in the metabolomic profiling studies. This highly characterized and diverse Bacteroidales culture collection with strain-resolved genomic and metabolomic analyses can serve as a resource to facilitate informed selection of strains for microbiome reconstitution.

## Introduction

Species of the Bacteroidales order are ubiquitous and abundant members of the human colonic microbiome with diverse impacts on host health. While heterogeneity amongst Bacteroidales species was recognized over 90 years ago,^1^ with subsequent molecular typing hinting at even greater taxonomic complexity,^2,3^ the breadth and depth of Bacteroidales diversity was not fully appreciated until the era of whole genome and deep metagenomic sequencing. Over the past two decades, comparisons of genome sequenced isolates and metagenome-assembled genomes have added new branches to the Bacteroidota phylogenetic tree, with renaming of some taxa and introduction of new names for various genera and species.^4–8^ Human gut Bacteroidales species are currently distributed to at least eight families. Of these families, Bacteroidaceae species are by far the best-studied with much less known about members of other families including Odoribacteriaceae, Rickenellaceae and Prevotellaceae.

Bacteroidales species encode many carbohydrate-active enzymes (CAZymes), including glycolytic enzymes, that enable them to break down and utilize a wide range of dietary polysaccharides.^9,10^ In the process of catabolizing polysaccharides, some Bacteroidales produce metabolites that include acetate, propionate and succinate, which impact gut mucosal integrity and immune defenses.^11–14^ However, studies also hint at species and strain level diversity in the production of various short chain fatty acids.^15,16^ In addition to dietary polysaccharides, some Bacteroidales species utilize host mucin glycans,^17^ which reduce the thickness of the mucus layer lining the gut mucosa, impacting immune regulation and antimicrobial defenses.^9,18^ The spectrum of Bacteroidales strains colonizing the host is impacted by diet, food preparation, and food components.^10^ While individual Bacteroidales species generally encode a core constellation of polysaccharide utilization loci (PULs) and metabolize a similar spectrum of polysaccharides,^17^ genetic exchange between species^19,20^ and loss of some functional PULs, such as those enabling host mucin glycan utilization, can lead to intraspecies diversity in utilizable substrates.^17^ We also know little about the glycolytic capabilities of many of the other families of gut Bacteroidales.

Bile acids are prevalent, cholesterol-derived steroidal molecules produced in the liver and introduced into the mammalian intestinal lumen as taurine and glycine conjugates that are sequentially modified by different members of the microbiota. Deconjugation and further modification of bile acids and other steroidal molecules by gut microbes is increasingly recognized to impact the mammalian host^21^ and is mediated, in part, by members of the Bacteroidales order.^22–25^ For example, a bile salt hydrolase synthesized by Bacteroides thetaiotaomicron VPI-5482 impacts host metabolic functions including weight gain and the metabolism of carbohydrates and lipids.^25^ A sulfotransferase produced by some Bacteroidales species can sulfonate cholesterol and a subset of secondary bile acids.^22,24^ Several Bacteroidales genera and species encode reductases and hydroxysteroid-dehydrogenases that modify secondary bile acids and produce variants that impact lymphocyte differentiation and may also inhibit enteric pathogens.^23^ A previous metabolomic study of 59 Bacteroidales strains of three families has been a useful resource to the field, and the current study includes additional Bacteroidales families and species and additional metabolites further highlighting the extreme species and strain level metabolic heterogeneity.

Another feature of gut Bacteroidales is their rapid evolution due to the exchange of DNA by conjugation. These mobile genetic elements encode numerous properties that affect strain fitness, including PUL that increase utilizable substrates,^19^ transport cofactors,^26^ Type VI secretion systems^27^ and bacteriocins^28,29^ that antagonize competing strains, along with protective immunity genes,^30^ and antibiotic resistance genes,^31^ to name a few. Although extensive conjugal DNA transfer between species within an individual’s gut microbiota has been documented,^32,33^ there are few studies that examine the degree to which Bacteroidales strains evolve and differentiate in the human gut due to DNA acquisitions.^34^ In addition, a better understanding of the phenotypes encoded on these mobile genetic elements and their impact on strain fitness and subsequent impacts on human host phenotypes becomes increasingly important as strains are selected as components of synthetic consortia for microbiome restoration.

Loss of microbiome diversity in a variety of settings is associated with adverse health outcomes and has led to studies of microbiome reconstitution with symbiotic gut bacterial strains or consortia, referred to as live biotherapeutic products (LBPs).^35^ As Bacteroidales strains are prevalent colonizers of the human intestine and generate metabolites that contribute to health and disease resistance, identifying those that produce beneficial metabolites is essential. Furthermore, identifying bacterial strains that successfully co-colonize with other species and support microbiome richness and diversity is also an important goal. In this study, we analyzed a large culture collection of diverse Bacteroidales strains of five families and 35 species isolated from healthy human donors to provide an order-wide analysis of their genomic, genetic and metabolomic properties. The substantial accessory genomes harbored by Bacteroidales strains encode a range of functions that, if defined, might be paired with the host recipient’s microbiome composition, metabolomic profile and underlying disease status, with the goal of optimizing the effectiveness and health impact of next-generation LBPs.

## Results

### Isolation, identification, and deduplication of Bacteroidales isolates

We collected stool samples from 21 healthy human donors, which were processed for 16S rRNA amplicon sequencing.^36^ Among the 21 donors, Bacteroidales represented 12.88% of the bacterial abundance on average (range 3.40% to 46.83%, median 8.32%) (Fig. S1). Fecal samples were diluted, and strains cultured on rich, non-selective media agar plates under anaerobic conditions. Single-colony isolates were sequenced using short-read whole genome sequencing (WGS). The culture collection was not designed to collect the diversity of Bacteroidales strains within each individual, but rather to build a diverse Bacteroidales strain bank for order-wide genomic, genotypic, and metabolomic analyses to facilitate future informed selection of consortia members for microbiome restoration.

To identify isolates of the Bacteroidales order, we performed BLAST searches using the longest 16S rRNA sequence from the WGS against the NCBI RefSeq 16S rRNA database or the Genome Taxonomy Database (GTDB). These analyses identified 408 strains of Bacteroidales (Dataset 1, Bacteroidales collection tab). To correctly speciate these isolates, we used 25 single-copy core genes (SCGs) included in the Anvi’o database^37^ (Dataset 1, single copy core genes tab). Multiple sequence alignment of the concatenated sequences of these 25 SCGs from the 408 Bacteroidales genomes and those of Bacteroidales type strains provided clear taxonomic assignments into 35 species of nine distinct genera (Fig 1A, Dataset 1, Bacteroidales collection tab).

**Figure 1.**
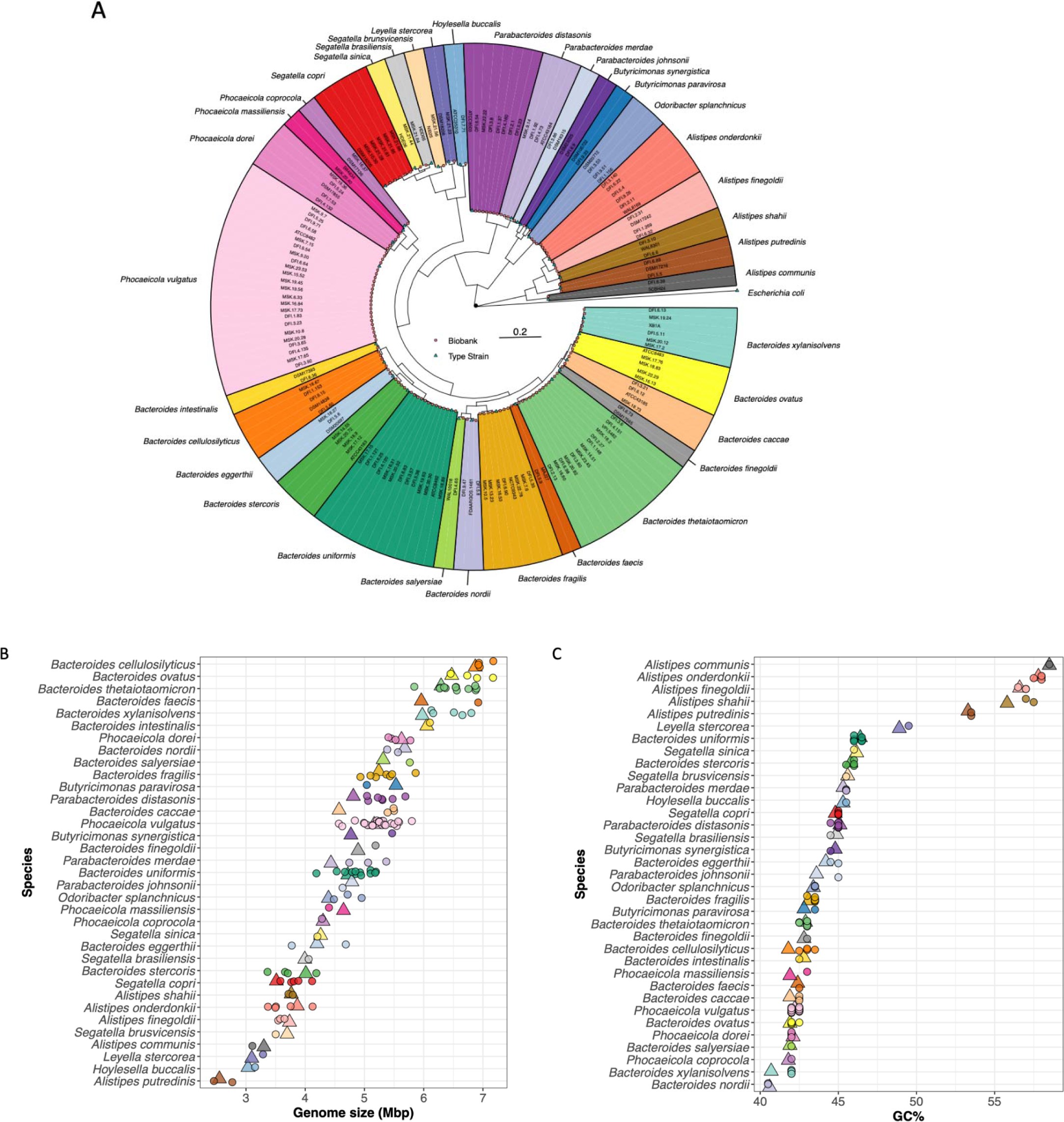
Phylogeny and genomic characterization of the 127 unique gut Bacteroidales isolates. A. Phylogenetic tree of unique gut Bacteroidales isolates and type strains based on concatenated single-copy core gene sequences. Escherichia coli serves as an outgroup. Scale bar indicates phylogenetic distance. 127 isolates (circles), type strains (triangles). B. Genomic size of human gut Bacteroidales isolates (circles) and type strains (triangles), arranged by decreasing average genome size of species from top to bottom. C. Genomic %GC content of human gut Bacteroidales isolates (circles) and type strains (triangles), arranged by decreasing average %GC content of species from top to bottom.

As many donor microbiotas contained multiple isolates of the same species, we sought to reduce redundancy by identifying clonal isolates. For this analysis, we performed pairwise comparisons among genomes of all strains of the same species using fastANI^38^ for Average Nucleotide Identity (ANI) and nucmer^39^ for genomic single nucleotide polymorphisms (SNPs) and insertion-deletions (InDels). Compared to isolates of the same species from different donors, those from the same donor typically have higher ANI and fewer genomic SNPs and InDels (Fig. S2A-C), with clear apparent cutoffs. We used the following criteria as thresholds for concluding isolates were of the same clonal lineage: ANI higher than 99.9%, total genomic SNPs fewer than 1000, and total genomic InDels fewer than 75. These analyses revealed many microbiomes where there were multiple distinct co-colonizing strains of the same species. To confirm that two strains of the same species from a microbiome are either the same or distinct clonal lineages, we performed pangenome analysis of these isolates using Anvi’o.^37^ This gene clustering-based approach clearly separated strains, as clonal isolates consistently have more than 95% identical gene clusters whereas isolates of different strains shared 80% or less (Fig. S3). Distinct co-colonizing strains of the same species were identified in nine microbiomes including DFI.3 where the four B. thetaiotaomicron isolates segregate into two distinct clonal lineages, and the five Phocaeicola vulgatus isolates segregate into three distinct clonal lineages (Dataset 1, de-duplication tab). Similarly, isolates from the MSK.21 microbiome were only of the Prevotellaceae family, including four different Segatella species,^5^ which is a characteristic of microbiomes from non-industrialized populations. In our strain collection, Ph. vulgatus was the most isolated species and also the species found most often to have two or more co-colonizing strains of distinct lineages (in six donor microbiota).

### Phylogeny of gut Bacteroidales based on curated single-copy core genes

We reduced the set of 408 strains by selecting only one strain of the same clonal lineage, which yielded 127 distinct clonal isolates (Dataset 1, de-duplication tab). Next, we built a phylogenetic tree of these 127 unique human gut Bacteroidales strains along with type strains of each of the 35 species using multiple sequence alignment of the concatenated sequences of the 25 SCGs (Fig. 1A, Dataset 1 single copy core genes tab). As expected, families (Bacteroidaceae, Prevotellaceae, Odoribacteraceae, Tannerellaceae and Rikenellaceae) and genera within these families (Bacteroides, Phocaeicola, Parabacteroides, Segatella, Leyella, Hoylesella, Butyricimonas, Odoribacter, and Alistipes) were well separated and formed monophyletic clades.

### Genome size and GC content of Bacteroidales isolates

We next analyzed genome size across Bacteroidales isolates (Fig. 1B). Genome sizes varied drastically between species of different families from as small as ∼2.8 Mbp (Alistipes putredinis) to >7.1 Mbp (Bacteroides cellulosilyticus and Bacteroides ovatus). In general, the largest genomes are from strains of Bacteroides known to have an extensive capability to digest host and dietary polysaccharides and that have the greatest number of carbohydrate active enzymes (CAZymes) and polysaccharide utilization loci (PUL). There was also considerable variation in the genome sizes within strains of the same species, the most dramatic example being genomes of Ph. vulgatus that ranged from ∼4.57 Mbp to ∼5.80 Mbp (Dataset 1, Bacteroidales collection tab). Intraspecies genome size discrepancies highlight the extensive horizontal gene transfer that occurs in these species, and the large pangenomes.

Analysis of %GC content revealed a wide range between species of this bacterial order, with Bacteroides nordii having the lowest at 40.5%, and A communis the highest at 58.5% (Dataset 1, Bacteroidales collection tab). There are typically narrow distributions of GC content within a species. Moreover, isolates within a genus tend to have similar trends in GC content, with Alistipes species having relatively higher GC content (53.5 - 58.5%) and Phocaeicola species having a narrow range and relatively lower GC content (42-43%) (Fig. 1C).

### Core genomes of Bacteroidales isolates

We annotated these Bacteroidales genomes using the Prokka pipeline with customization (see Methods).^40^ The number of coding regions varied from 2167 to 5671, due to drastic differences in genome size, with an average of 75% of all putative CDS being annotated (range: 65% to 86%). To investigate the core genomes across taxonomic levels, we performed annotation-agnostic clustering of all CDS of the 127 unique Bacteroidales isolates and their type strains. 247 gene clusters were shared across all isolates (order-level core genome)(Fig. 2). At the family level, the core genome size increased to 979 gene clusters for Bacteroidaceae, 842 for Prevotellaceae and 1366 for Odoribacteraceae. Other families included only one genus, so family level analysis was not possible. Genus level analysis was included for all genera with more than one species and further increased the size of the core genomes: 1175 gene clusters for Bacteroides, 1438 for Phocaeicola, 1862 for Parabacteroides, 2045 for Butyricomonas, 992 for Alistipes, and 1366 for Segatella. Strains of the same species shared on average about 69.6% of core genomes (median 69.5%, range 52.5%-89.8%) (Fig 2).

**Figure 2.**
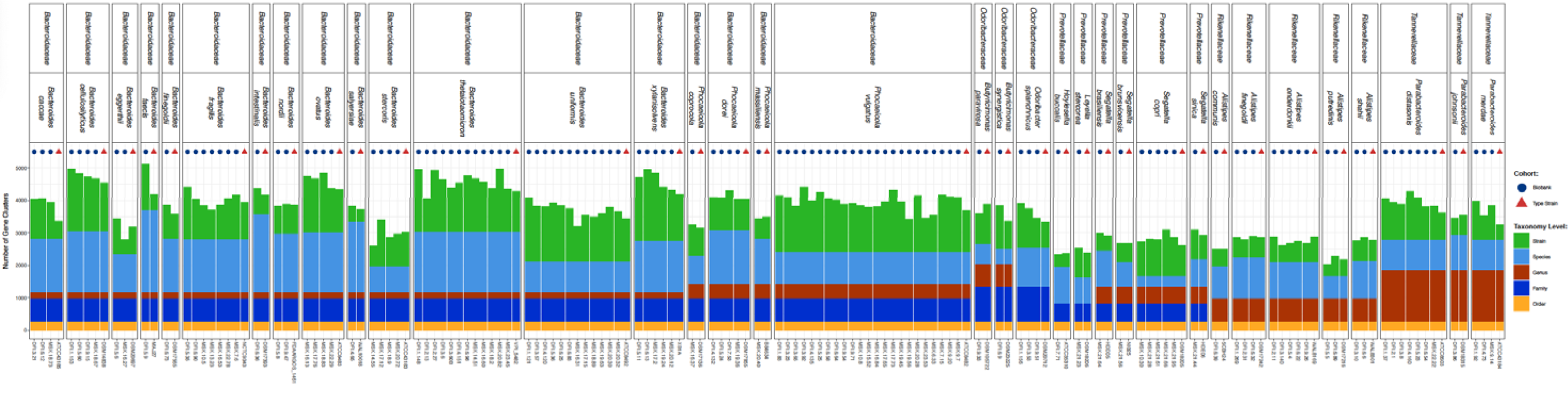
Core genomes of Bacteroidales isolates at various taxonomic levels. The number of core gene clusters shared by isolates grouped at different taxonomic ranks is plotted. Color indicates core gene clusters in different taxonomic ranks. Bacteroidales collection isolates (circles) or type strains (triangles).

### Diversity of Bacteroidales based on genetic repertoire

To further elucidate the genomic diversity among human gut-derived Bacteroidales, we performed Uniform Manifold Approximation and Projection (UMAP)^41^ based on the presence or absence of unique gene clusters from their whole-genome protein coding sequences (Fig. 3, Fig. S4). We found that different families and genera were well separated in UMAP, and in most cases, isolates of the same species and their corresponding type strains clustered together in UMAP and were distinguished from other species (Fig. 3, Fig. S4). P. vulgatus isolates occupied a larger area compared to other species, and some P. vulgatus isolates were closer to type strains of other Phocaeicola species than to the P. vulgatus type strain (Fig. 3 and Fig. S4) with strain distances similar to those of other species of a genus.

**Figure 3.**
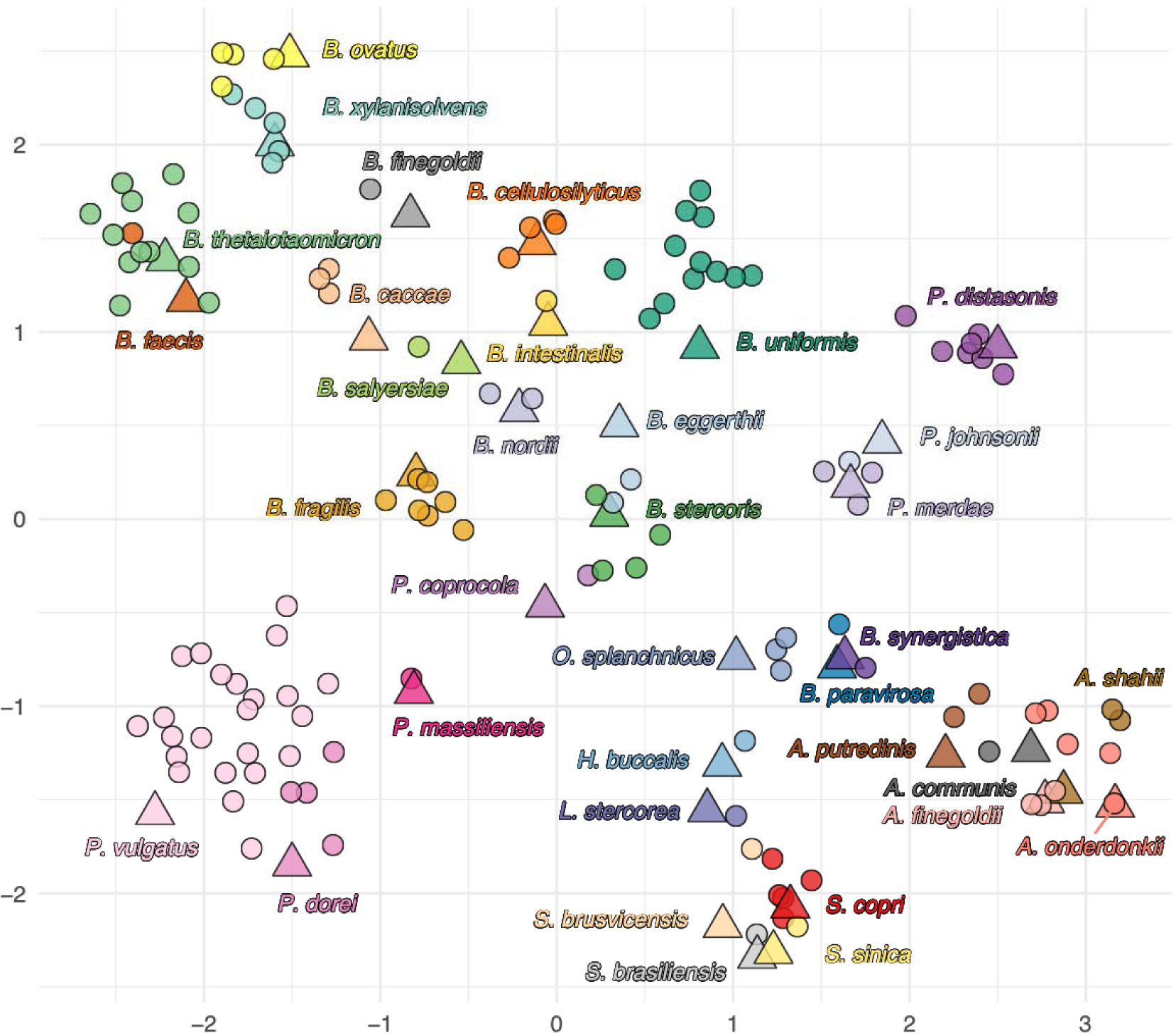
UMAP analysis of the genomic diversity of Bacteroidales isolates. UMAP plot based on the presence or absence of gene clusters in whole-genome protein coding sequences of all Bacteroidales human isolates and type strains in our collection. Bacteroidales collection isolates (circles) or type strains (triangles).

### Distribution of CAZymes and PULs among human gut Bacteroidales

The distribution of CAZymes and PUL has been extensively characterized in gut Bacteroidales species as these features indicate a strain’s glycan utilizing capabilities. To determine the distribution of CAZymes in this Bacteroidales culture collection, we used Hidden Markov Models from dbCAN database^42^ to search for CAZymes and categorized them according to the CAZy database.^43^ We detected 305 CAZy families/subfamilies among Bacteroidales isolates with marked variation in the total number of CAZymes in different species (Fig. 4A and Fig. S5A). Most identified CAZymes are of the Glycoside Hydrolases family (GH), followed by Glycosyl Transferases (GT), Polysaccharide Lyases (PL), Carbohydrate Esterases and Carbohydrate-Binding Modules (Fig. 4A). Bacteroides cellulosilyticus contained the largest numbers of total and unique CAZy families/subfamilies present, whereas Alistipes putredinis contained the least (numbers of total CAZy families/subfamilies in Fig. 4A, and numbers of unique CAZy families/subfamilies in S5B). There are trends in the categories of CAZymes within a genus, with Bacteroidaceae species typically containing the most glycolytic enzymes (GH and PL). The number of CAZymes present in a species also correlated with the diversity of their CAZymes measured by Inverse Simpson Index (Fig. S5A). Glycosyl Transferases are CAZymes that are involved in the synthesis of glycans present as surface molecules (capsular polysaccharides, extracellular polysaccharides, surface layers) or components of Bacteroidales glycoproteins.^44,45^ There is also extreme diversity in the number of GT between species with Alistipes putredinis encoding only 10 GT and some P. dorei strains encoding as many as 80. These differences indicate that that the ability to synthesize numerous phase variable surface polysaccharides, which is a trait of the gut Tannerellaceae and Bacteroidaceae,^46^ is not a trait shared by all gut Bacteroidales.

**Figure 4.**
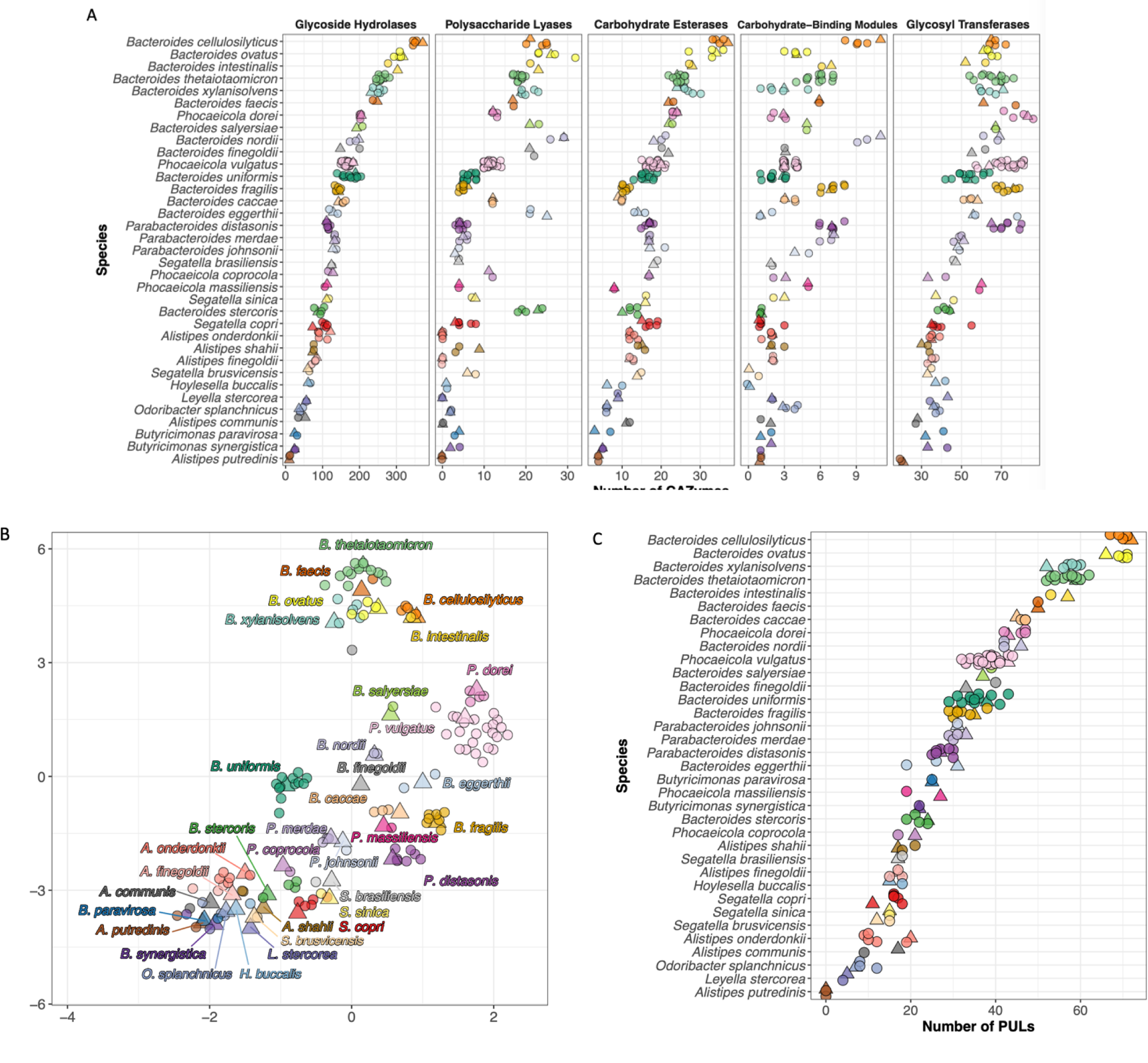
CAZyme and PUL analyses of Bacteroidales isolates. A. Total numbers of CAZymes in each Bacteroidales isolate, arranged by species from most (top) to least (bottom). CAZyme families are categorized into five major classes according to CAZy database, each panel represents one class. B. UMAP plot based on the presence or absence of CAZyme families in all Bacteroidales human isolates and type strains used here. Color indicates species identity. Bacteroidales collection isolates (circles) or type strains (triangles). C. Number of PULs in all Bacteroidales isolates and type strains, ranked by their average PUL numbers from most (top) to least (bottom).

UMAP analysis based on the presence or absence of CAZyme families revealed that isolates of the same species tended to cluster together (Fig. 4B). Bacteroides thetaiotaomicron, B. xylanisolvens, B. ovatus, B. intestinalis and B. cellulosilyticus— species with the most CAZymes— are distant from other clusters, suggesting they possess distinct CAZyme profiles (Fig. 4B). Alistipes and Odoribacteraceae species, which encoded the fewest CAZymes, cluster together (Fig. 4B). Analysis of the CAZyme UMAP based on donor showed that isolates from the same individual are scattered around the UMAP (Fig. S5C) supporting that different Bacteroidales species occupy distinct nutritional niches in the microbiome.

Exploitative competition, or the ability to compete for nutrients, often limits co-colonization of strains of the same species in a microbiome. Successful co-colonization may require utilization of distinct glycan sources. We analyzed the CAZymes present in the three distinct P. vulgatus strains that co-colonized donor DFI.3 (DFI.3.23, DFI.3.65 and DFI.3.92) and found strain differences in the CAZyme families. GH99 are endo-α-mannosidases that cleave mannoside linkages in N-linked glycan chains.^47^ GH99 is present in single copy in DFI.3.65, with two copies in DFI.3.92, but absent in DFI.3.23. Two different GH families, GH10 and GH67 are uniquely present in DFI.3.92. GH10 members are endoxylanases^48^ and GH67 members are xylan α- glucuronidases,^49,50^ some of which are upregulated in Bacteroides species upon culturing with xylan.^50^ These findings suggest that P. vulgatus strains within the same donor have unique capabilities to utilize different dietary/host components, exploiting distinct nutritional niches in the same microbiome. We further analyzed the CAZyme profiles of all strains of the same species of distinct clonal lineages co-colonizing individuals and similarly found extensive differences, likely accounting for their ability to co-colonize the same host (Dataset 2). Interestingly, among the CAZyme families that are discordantly present within each comparison, GH46 is the most frequent. GH46 encodes a predicted chitosanase not previously characterized in Bacteroidales. Other frequently discordant GH families include GH117, a neoagaro- oligosaccharide hydrolase family in an agarose metabolic pathway shown to be acquired by a B. uniformis strain,^51^ and GH96, an α-agarase family previously uncharacterized in Bacteroidales.

In Bacteroidales, the majority of glycolytic enzymes are encoded in PULs. Although PULs often contain genes encoding surface and periplasmic glycolytic enzymes, as well as regulatory genes,^52^ the minimum requirement for a locus to be considered a PUL is a set of susC/susD genes, encoding the outer membrane/surface glycan import machinery, and a gene encoding a glycolytic enzyme.^52^ We used PULpy^53^ to identify PULs in our Bacteroidales strain collection. Bacteroides cellulosilyticus strains contained the most PULs on average (67-72 PULs), followed closely by B. ovatus isolates (66-71 PULs)(Fig. 4C), consistent with these strains containing the most CAZymes (Fig. 4A, Fig. S5B). Alistipes putredinis contained only one susC/susD pair without a nearby GH or PL encoding gene. In general, Bacteroides species contained more PULs whereas Alistipes and Odoribacter species contain fewer (Fig. 4C), suggesting limited use of dietary and host glycans by these latter species.

### Bile acid metabolism capacities among human gut Bacteroidales

Gut bacteria produce enzymes that modify host-generated primary bile acids into secondary bile acids, which have numerous effects on both the host and the microbiota. Many Bacteroidales produce bile salt hydrolases (BSH) that deconjugate the taurine or glycine moiety which enables further modification by other microbiota-encoded enzymes.^21,25,54,55^ Using three distinct BSH sequences as queries, we found that most gut Bacteroidales of our collection have at least one BSH ortholog with the exception of species of Prevotellaceae, most Odoribacter splanchincus and the single B. finegoldii isolate (Dataset 3). Therefore, this important deconjugation reaction is performed by most gut Bacteroidales. A recent study showed that centenarians have increased levels of fecal lithocholic acid (LCA) including iso-LCA, allo-LCA, 3- oxo-LCA, 3-oxoallo-LCA and isoallo-LCA.^23^ We searched our Bacteroidales collection for 5α- reductase (5-AR) and 5β-reductase (5-BR) that produce 3-oxoallo-LCA and 3-oxo-LCA, respectively (Dataset 3).^23^ Of the 127 strains, the majority contain orthologs of both 5AR and 5BR with the exception of all Prevotellaceae, Bacteroides nordii, O. splanchinicus and one outlier B. uniformis strain, all of which lacked both proteins. In addition, Parabacteroides distasonis isolates contain only an ortholog of 5BR, but not 5AR. The genes encoding 5AR and 5BR are adjacent as previously described (Dataset 3).^23^ Lastly, we searched for hydroxysteroid dehydrogenases (HSDH). With the exception of the Prevotellaceae, Bacteroides nordii, O. spanchinicus and the same outlier B. uniformis strain, most Bacteroidales contain at least one ortholog of 3βHSDH, which produces isoalloLCA from 3-oxoallo-LCA. These genes are adjacent to the 5AR and or 5BR genes as described.^23^ 7αHSDH is present in only some Bacteroides species and one O. splanchinicus isolate (Dataset 3). Collectively, these data indicate that most Bacteroidales strains encode enzymes that can modify bile acids to produce a variety of secondary bile acids, with species and strain level diversity. It remains unclear if the Prevotellaceae have bile acid conversion capacities, but our data show that they do not have close ortholog to any of the known Bacteroidales proteins with these functions.

### Metabolome of Human Gut Bacteroidales

Human gut Bacteroidales produce a wide range of metabolites that affect other microbes and impact host phenotypes.^11–13,56,57^ We performed metabolomic analyses of culture supernatants of Bacteroidales strains grown in BHIS medium. We quantified the short chain fatty acids (SCFAs) acetate, propionate, butyrate and the dicarboxylic acid succinate. In addition, a panel of 29 metabolites was analyzed semiquantitatively, enabling us to compare relative metabolite amounts to baseline levels present in the medium to determine production versus consumption. The metabolomes of 111 strains that grew well in BHIS were included.

Bacteroidales strains produce acetate, succinate and propionate, with some exceptions. For example, B. stercoris MSK.20.72 did not produce acetate, produced only small amounts of propionate and consumed succinate. Butyrate was produced by Odoribacteraceae isolates (two Odoribacter splanchnicus and one Butyrimonas synergistica) (Fig. 5A, Fig. S6A). These O. splanchnicus isolates contain both Acetyl-CoA-Type and Lysine-Type complete butyrate production pathways. The genomes of both Alistipes putredinis strains also encode complete pathways for butyrate production, however, when grown in BHIS, they produced only very low amounts of butyrate (0.055-0.14 mM compared to 0.54-18 mM from O. splanchnicus) (Fig. S6A). Most notable in these analyses are numerous examples of extreme intra-species diversity in the production of these metabolic end products.

**Figure 5.**
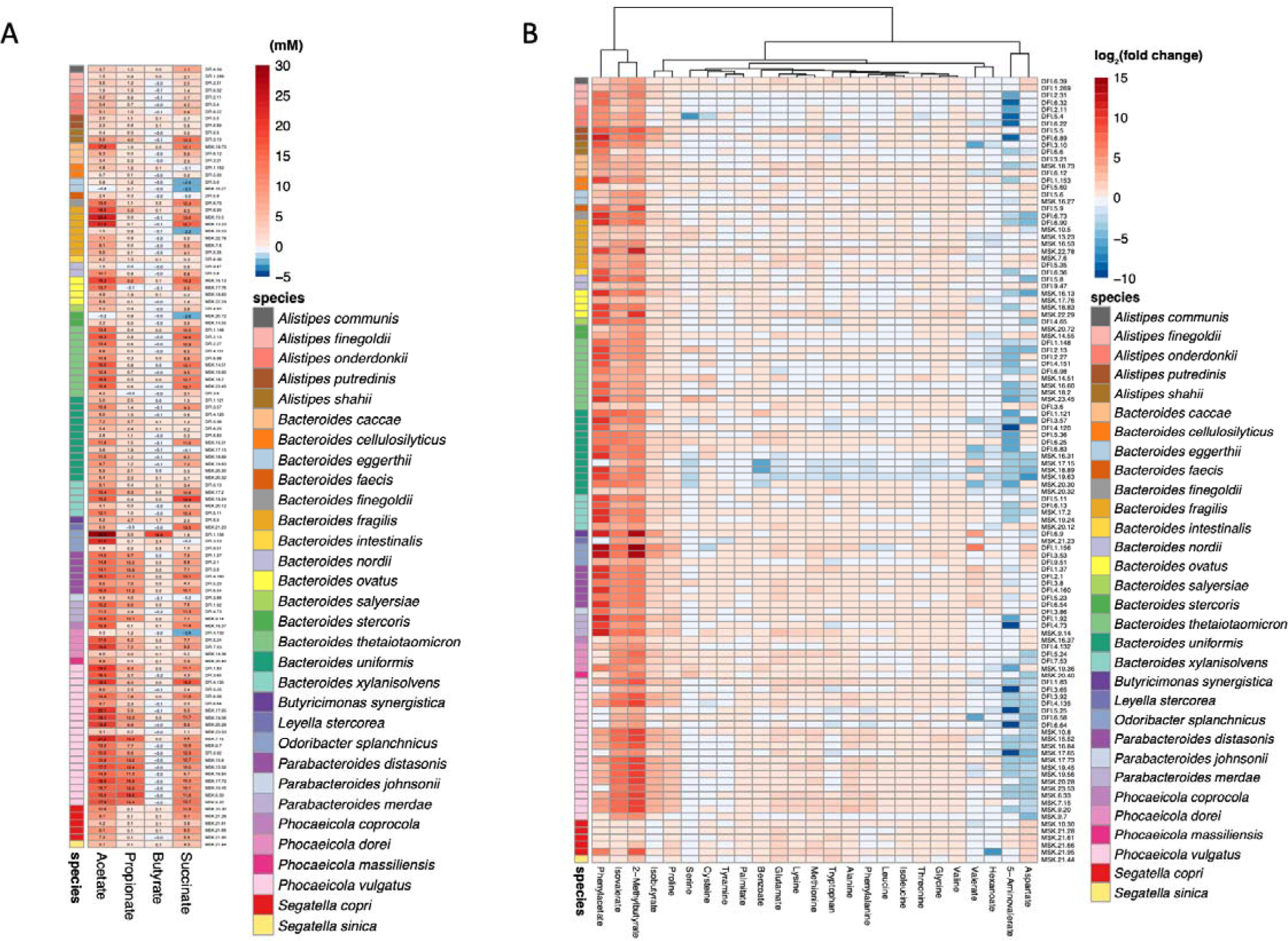
Metabolomic profiles of Bacteroidales isolates. A. Quantitative metabolomics on SCFAs among Bacteroidales isolates. Each column represents one SCFA. Numbers in wells indicate relative concentration changes in mM compared to BHIS medium. Color squares on the left indicate species identity. B. Semi-quantitative metabolomics on 25 metabolites among Bacteroidales isolates. Each column represents one metabolite. Columns are grouped by hierarchical clustering. Color scale indicates log_2_(fold change) of metabolite concentration relative to BHIS medium. Color squares on the left indicate species identity.

The relative production versus consumption of metabolites, such as branched-chain and aryl acids, amino acids, medium-chain fatty acids, among Bacteroidales strains was also determined and demonstrated, for some metabolites, significant inter- and intra-species variation (Fig. 5B).

One of the most variably produced metabolites in our panel is phenylacetate. Gut microbiota-derived phenylacetate is the critical precursor for host production of phenylacetylglutamine, which is associated with both development of atherosclerotic cardiovascular disease, major adverse cardiovascular events, and stroke risk.^58–60^ Phenylacetate is reported to be produced from phenylalanine via two different pathways by some gut microbes, including Bacteroides species,^57^ and involves specific oxidoreductases. We observed that in some species, such as B. uniformis and B. thetaiotaomicron, the numbers of FAD-dependent oxidoreductases encoded in their genomes correlates with phenylacetate production (Fig. S6B, S6C).

Hierarchical clustering of metabolomic profiles did not precisely follow phylogeny (Fig. S6D), consistent with previous findings.^16^ Procrustean Approach to Cophylogeny (PACo) test did not detect a significant correlation between phylogeny and metabolomic profiles (P = 0.129) (Fig. S6E). Thus, our metabolomic results suggest that while genomic analyses can in many cases determine whether production or consumption of specific metabolites occurs, inter- and intra-species diversity of metabolomes cannot, at this time, be fully explained by genomic composition, highlighting the need for broader and deeper analyses of bacterial metabolite production or consumption and gene transcription under varying culture conditions.

### Analysis of genes encoding antagonistic proteins/machinery

Bacteroidales species, like most species that live in communities, produce many antagonistic weapons to thwart competitors. To date, five distinct antibacterial toxin families (BSAPs, BfUbb, BcpT, bacteroidetocins, and Fab1) and three distinct Type VI secretion systems (T6SSs) architectures (GA1, GA2 and GA3) have been identified. Many of these antibacterial toxins have been shown to limit competing species in the mammalian gut.^61–66^ We queried the genomes for genes encoding BSAP-1 – BSAP-4,^61,67,68^ involved in intra-species antagonism in B. fragilis, B. uniformis, Ph. vulgatus and Ph. dorei; BcpT,^28^ which is restricted largely to Ph. vulgatus and Ph. dorei; bacteroidetocin A and B, which are widely distributed among gut Bacteroidaceae;^29^ and Fab1^62^ and BfUbb,^69^ which are restricted to B. fragilis. As expected, a subset of strains of the expected species contains the genes encoding each of these toxins with the exception of BfUbb, which was not encoded in any of the 7 B. fragilis genomes in our collection, consistent with a previous reported prevalence of 14% among sequenced B. fragilis strains.^69^ There were two other notable findings from this analysis. The first is that bcpT and the 9.1 kb plasmid that encodes it were found not only in Ph. vulgatus and Ph. dorei as expected, but also in B. caccae, a species not previously shown to harbor this mobile genetic element. The second interesting finding is that BcpT and Bd-B, both encoded on mobile genetic elements (MGEs), are present in some isolates of the same clonal lineage but not all (see below), indicating that these mobile elements were acquired or lost by some strains in those individuals’ gut microbiome.

To investigate T6SS distribution, we performed BLAST searches using concatenated T6SS consensus sequences of each of the three GA types and the five GA2 subtypes.^33,70^ We detected at least one T6SS in most genera except Butyricimonas, Segatella and Hoylesella, which extends Bacteroidales species containing T6SS loci to Odoribacter and Alistipes. However, in the Odoribacter spanchnicus isolates of DFI.1, 14 genes the GA1 T6SS region are present in the reverse orientation, resulting in the disruption tssD and tssC for which partial genes are contained on each side of the inverted region (Fig S6A) leading to a non-functional T6SS. Of the 127 unique isolates analyzed, GA1 is present in 20 genomes, GA2b in 14 genomes, GA2a in 5 genomes, GA2c in 1 genome, and GA3 in 4 genomes. GA1 and GA2 T6SSs loci are present on integrative and conjugative elements (ICE) and we found them to be variably present in strains of the same clonal lineage (see below), demonstrating acquisition of these antagonistic weapons by strains in the microbiota of these human subjects. Our previous analyses had revealed GA2 ICE exclusion wherein we had not previously identified stains with both a GA2 and GA3 T6SS, and rarely found isolates containing both a GA2 and GA1 T6SS loci. Of those genomes that did contain both GA1 and GA2 ICE, we previously identified a defect in one of the T6SS loci or in a gene adjacent to the GA1 T6SS locus designated mhgA.^33^ Three strains in our collection, B. uniformis DFI.1.121, B. thetaiotaomicron DFI.1.148 and B. thetaiotaomicron DFI.2.27, contain both GA2a and GA1 T6SS loci in their genomes (Dataset 1, T6SS tab), and in each, the mghA gene is similarly interrupted by the insertion of a putative transposase-like element encoding a reverse transcriptase (Fig. S6B). MhgA is a protein containing both an N - adenine methylase domain and an SNF2 helicase domain, which is similar to protective genes in the phage-defense system (DISARM).^71^ The most surprising result of these analyses was the identification of a B. fragilis strains with all three GA types (GA1, GA2a, and GA3) (Fig. S7; Dataset 1, T6SS tab). In this strain, the mhgA gene of the GA1 ICE was also interrupted, but with a different transposable element (Fig S7).

### Horizontal DNA transfer between co-resident species

We and others have previously demonstrated extensive transfer of conjugative elements between Bacteroidales species in a person’s gut microbiota.^27,32,72^ However, most of these transfers were detected within and between species of Bacteroidaceae and Parabacteroides. As our culture collection contains diverse Bacteroidales species of five different families, we had the opportunity to identify DNA transfer among these diverse families. We used previously established criteria of 99.99% DNA identity over a minimum of 5 kbp to detect elements transferred between species within an ecosystem. Using the entire set of 408 isolate genomes, we detected extensive transfer of DNA between co-resident species, which included mobile and conjugative plasmids, integrative and conjugative elements (ICE), lysogenic phage and other mobile genetic elements (Dataset 4). As mentioned above, these transfers included the GA1 and GA2 T6SS loci that are contained on ICEs. In the gut microbiome of donor DFI.1, a GA1 ICE transferred among seven different species of three families: Bacteroidacae, Tannerellaceae, and Rickenellaceae. We also detected transfer of other genetic elements between Bacteroidaceae and Odoribacter, Butyricomonas and Alistipes (Dataset 4). The community of donor MSK.21 contained five different Prevotellaceae species, from which we detected DNA transfers among Leyella stercorea and Segatella brunsvicensis. Figure 6A illustrates five different elements that were transferred between different species in the microbiota of donor DFI.9. The full list of these genetic regions and their putative annotations are included for each ecosystem (Dataset 4). These analyses also highlight the ubiquity of lysogenic phage transferring between species in the human gut. We identified 20 phages that transferred between Bacteroidales species in this collection — one as large as 102 kb — and all of them belong to the Cauldoviricetes group which are contractile tail phages. In donor DFI.1, B. cellulosilyticus, O. splanchnicus, and Parabacteroides merdae, comprising three different Bacteroidales families, all contained the identical prophage in their genomes, demonstrating a broad host range.

**Figure 6.**
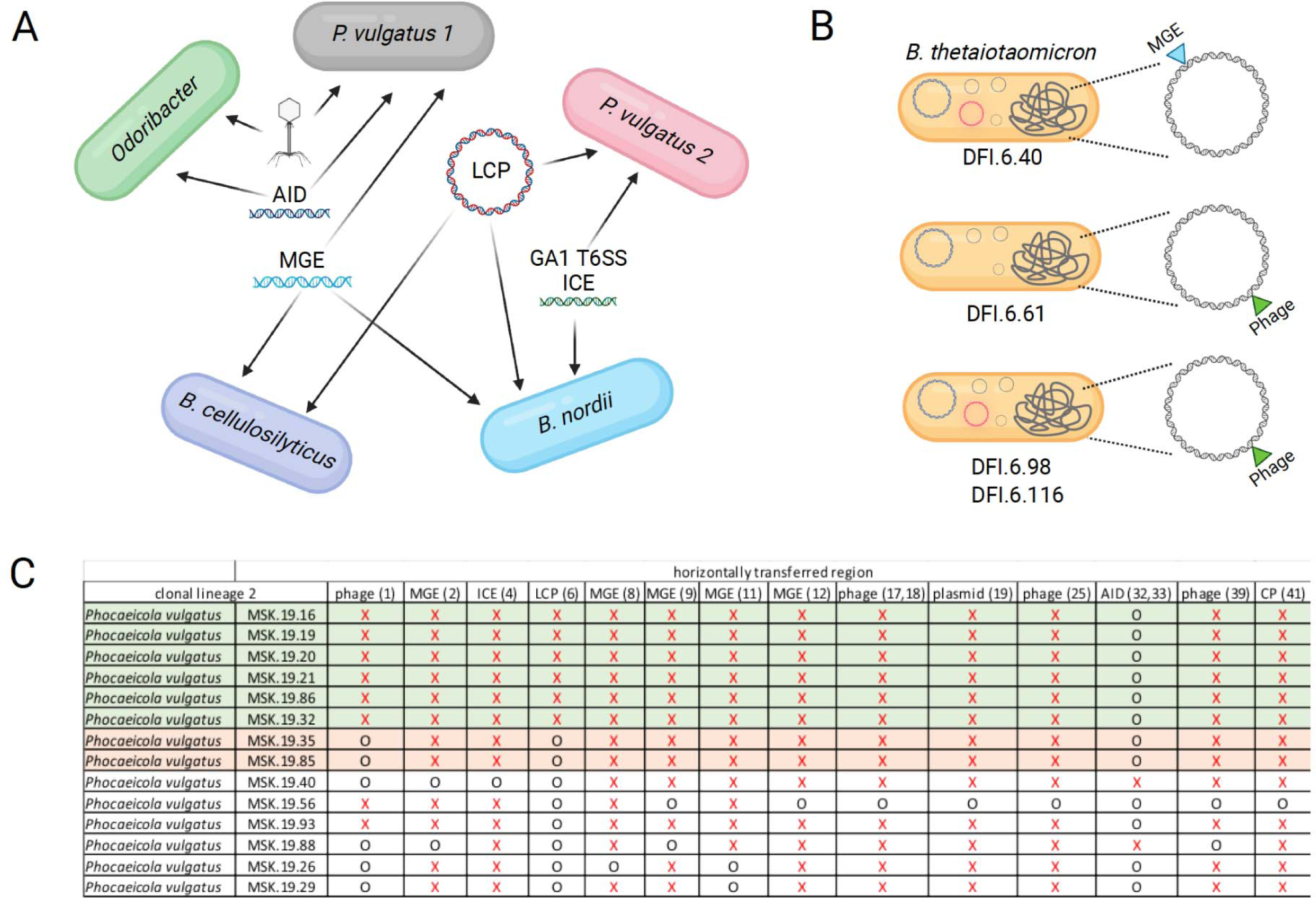
Strain evolution in the human gut microbiota due to DNA transfer events. A. Phage/genetic elements that transferred between the various species of the DFI.9 microbiome. B. Divergence and evolution due to DNA acquisitions of four B. thetaiotaomicron isolates of DFI.6 of the same clonal lineage. Sites of insertion of the elements in each of the genomes is shown. The MGE is 29,213 bp, the phage is 10,545 bp and all strain contain plasmids of 98,230 bp, 4,971 bp, 4,148 bp and 2,750 bp, and three of the four strains also contain a 44,031 bp plasmid shown in pink. C. Extensive strain divergence of a clonal lineage of P. vulgatus of ecosystem MSK.19 due to differential DNA acquisitions/losses. Eight distinct strains are differentiated based on the 14 transferred elements listed across the top. Strains that are the same in regard to the presence or absence of these elements are shaded with the same color. Refer to Dataset 4 for details regarding the genes of these regions. Red X indicates presence of the element and black O indicates the element is absent in that strain. Number in parenthesis refers to the region listed in Dataset 4, and this Dataset also contains details of the genes of these regions. AID – acquired interbacterial defense element. LCP – large conjugative plasmid.

We also detected numerous plasmids transferred among species. This size cutoff imposed in this analysis excluded pBI143, a ubiquitous mobile plasmid of ∼2. 8 kb that was recently described as one of the most abundant genetic elements of the human gut microbiome.^73^ We found pBI143 in strains of 16 of the 21 ecosystems and 40 of the 127 clonal lineages, including 14 species. In addition, we found another small plasmid of ∼5 kbp that was present in strains of seven communities. The largest plasmid detected was 105 kbp and is a variant of pMMCAT, a large conjugative plasmid carrying fimbriae genes and an extracellular polysaccharide biosynthesis locus.^33^ We previously found pMMCAT to be one of the most highly transferred elements with a conserved architecture.^33^ In our collection, we found evidence of transfer of pMMCAT-like plasmids in at least six communities.

### Strain evolution within the gut microbiota

Although we know that Bacteroidales strains evolve in the human gut due to mutation and DNA transfers mediated by conjugation and phage,^19,27–32,34,72,74^ tracking the evolution of a strain due to these DNA acquisitions or losses requires the analysis of numerous clonal strains of the same lineage from an individual. During our strain de-duplication process, we grouped strains into “clonal” isolates, i.e., those likely derived from a single strain acquisition by an individual. Seventy of the 127 de-duplicated strains contained more than one isolate of the same clonal group (Dataset 1, de-duplication tab). Within these clonal groups we searched for DNA regions of at least 5 kbp that were present in only a subset of clonal strains, but not all. These analyses revealed, in many cases, the divergence and evolution of strains due to the presence of various MGE including plasmids, ICE, and lysogenic phage. In fact, 45 of these 70 clonal lineages revealed strain evolution and divergence based on the presence or absence of MGE (Dataset 5). In Figure 6B we illustrate the genetic differences in the four B. thetaiotaomicron clonal isolates of DFI.6, showing the location of a 29,213 bp MGE that inserted just downstream of dnaK in one strain, and the insertion of a 10,545 bp phage in a gene of unknown function (DUF3822) in three of the four strains. All four strains also contain four plasmids, and three of the four strains contain an additional plasmid of 44,031 bp. Even greater divergence was detected in Ph. vulgatus isolates of clonal lineage 2 of the gut community of donor MSK.19, where we detected eight different Ph. vulgatus “strains” that evolved from a single ancestral strain acquired by that individual (Fig 6C). In addition to prophage, the genetic elements variably present in these eight evolved strains encode proteins involved in nutrient acquisition, restriction and modification enzymes, fimbriae, proteases, and antibacterial defense proteins to name a few (Dataset 5). In other clonal lineages, strains variably contain elements encoding T6SSs, antibacterial proteins and peptides, and proteins conferring antibiotic resistance. These data reveal the rapid and extensive evolution and divergence of clonal strains of Bacteroidales in the human gut due to horizontal gene transfer and the numerous phenotypic capabilities that likely accompany these acquisitions.

## Discussion

Species belonging to the Bacteroidales order constitute a major fraction of the lower intestinal tract microbiota and contribute to the host’s physiology and immune defenses. Here, we characterize a large and diverse Bacteroidales culture collection combining genome sequencing and extensive genomic and genetic analyses, with metabolomic analyses of individual Bacteroidales isolates. To our knowledge, this is the first Bacteroidales stain collection that is fully sequenced, characterized at the genomic, genetic, and metabolomic levels, with individual strains available to the research community.

The characterization of this strain bank has reinforced some properties of this order of bacteria and revealed new findings. For example, it was shown, first by culture-based analyses^75^ and more recently by single cell analyses^34^ that two or more strains of distinct clonal lineages of a species can co-colonize the human gut. The genetic and phenotypic differences that support such a co-occurrence have not been well-described. Our strain bank has captured numerous gut microbiomes in which multiple strains of distinct lineages of species were isolated, including multiple instances of co-colonizing Ph. vulgatus, B. thetaiotaomicron, and B. uniformis strains. In all cases, we find that co-colonizing strains have distinct accessory genomes including genes for PULs and other genes that likely support their co-colonization. Among our collection, Ph. vulgatus is also notable in having the greatest variation in genome size, size of the accessory genome, and wide distribution on the UMAP plot. Although some of this variation may be because our collection contains more Ph. vulgatus than other species, this diversity likely contributes to the frequent presence of multiple Ph. vulgatus strains of distinct lineages in a single gut ecosystem.

Not only do distinct lineages of a species co-colonize, but so to do numerous isolates of the same clonal lineage that have diverged due to the acquisition or loss of genetic elements. The properties conferred by the genetic cargo of these MGE are beginning to be characterized and include the utilization of dietary polysaccharides,^19,20^, cofactor import,^26^ antagonistic weapons,^27–29^ protection from T6SS mediated antagonism,^30^ transcriptional regulators that influence expression of chromosomal genes,^76^ biofilm formation, and antibiotic resistance.^77,78^ It can be anticipated that introduction of new Bacteroidales strains for therapeutic purposes will evolve differently in each ecosystem into which they are introduced and will also shape the Bacteroidales strains preexisting in or acquired by each ecosystem.

Metabolomic profiling of Bacteroidales isolates revealed significant inter- and intra-species diversity. While butyrate production was restricted to a small subset of species encoding genes for a complete butyrate production pathway, it remains unclear whether these symbiotic strains are significant contributors to the overall butyrate pool in the colon. Along similar lines, while we identified a correlation between phenylacetate production levels with the number of FAD-dependent oxidoreductases encoded in Bacteroidales genomes, further study is required to determine the impact of these strains and the production of phenylacetate on the host. Our finding that the metabolomic output of strains did not consistently align with phylogeny, though interesting, likely results from the limited media conditions under which we analyzed strain metabolomes. It is likely that media compositions that more closely reflect the in vivo environment that Bacteroidales strains encounter, including the presence of other microbial species, would provide insights into the regulation of metabolite production.

Bacteroidales species have been associated with disease susceptibility, on the one hand, and health benefits, on the other hand. The broad range of genomic and metabolomic features associated with individual strains within a given Bacteroidales species suggests that strains to be used for therapeutic purposes should be selected carefully.^79–81^ Our strain-resolution Bacteroidales repository provides detailed description and characterization of these human gut isolates, which are available to the scientific community (both in NCBI RefSeq database and at the DFI Symbiotic Biobank website https://dfi.cri.uchicago.edu/biobank/) and will facilitate development of targeted therapeutics and defined synthetic microbial consortia.^35,82^

## Acknowledgment

We thank members of the E.G.P. laboratory who have provided valuable feedback on this manuscript. Z.J.Z was supported by The GI Research Foundation early career grant. This work was supported by US National Institutes of Health grants P01 CA023766 to E.G.P. and by the Duchossois Family Institute of the University of Chicago. We also acknowledge the important role of the late Eric Littmann in initiating this study of the Bacteroidales order.

## Supplementary Figure Legends

**Figure S1.**
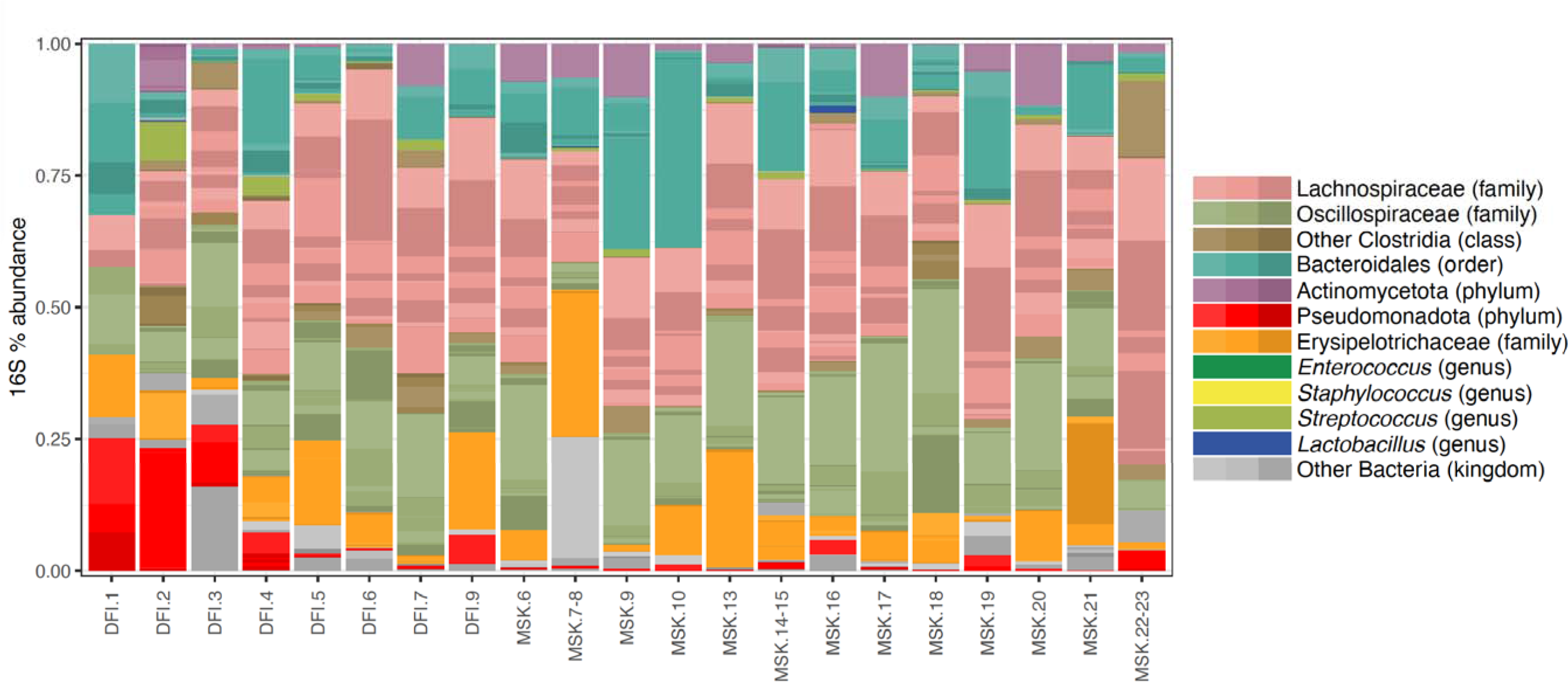
Gut microbial composition of human donors. Bar plot of relative abundance of bacterial taxa according to 16S rRNA amplicon sequencing. Each column is a human donor fecal sample.

**Figure S2.**
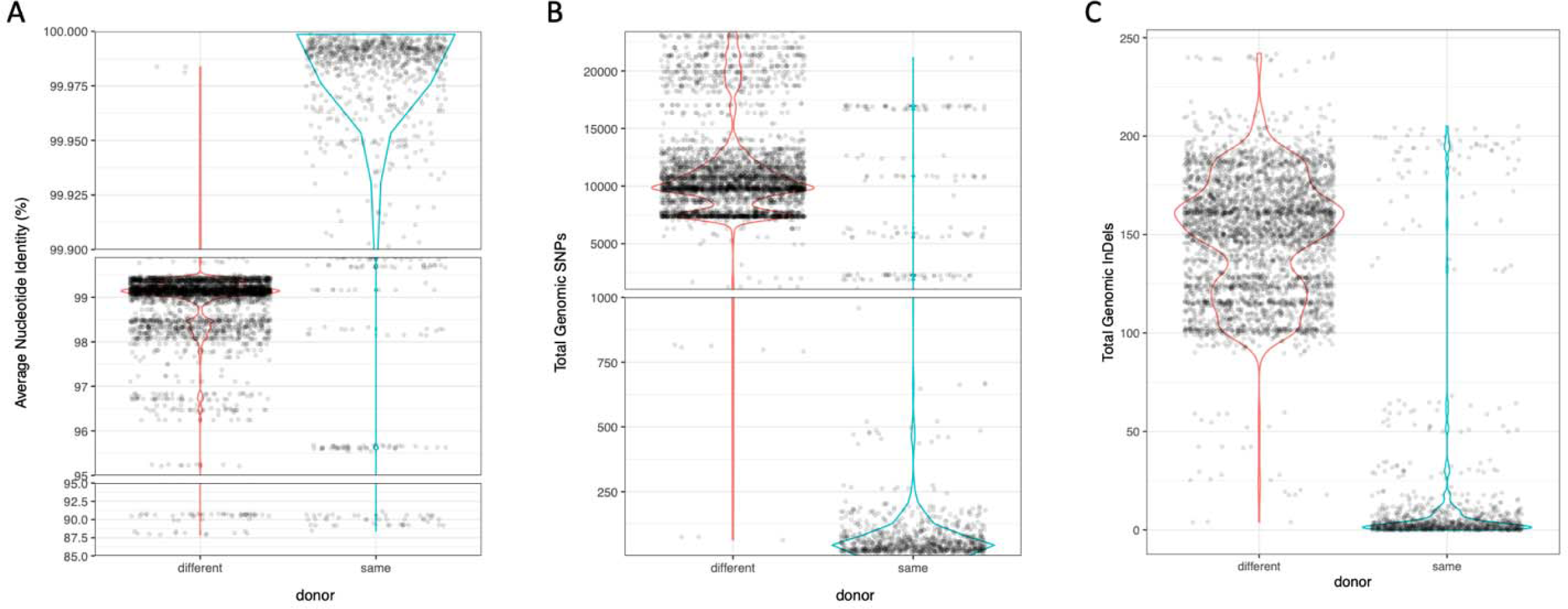
De-duplication of Bacteroidales isolates. Violin plots showing pairwise differences of Average Nucleotide Identity (A), total genomic SNPs (B) and total genomic insertions-deletions (C) between all isolates of same species. Each dot represents one pairwise comparison of two isolates of same species.

**Figure S3.**
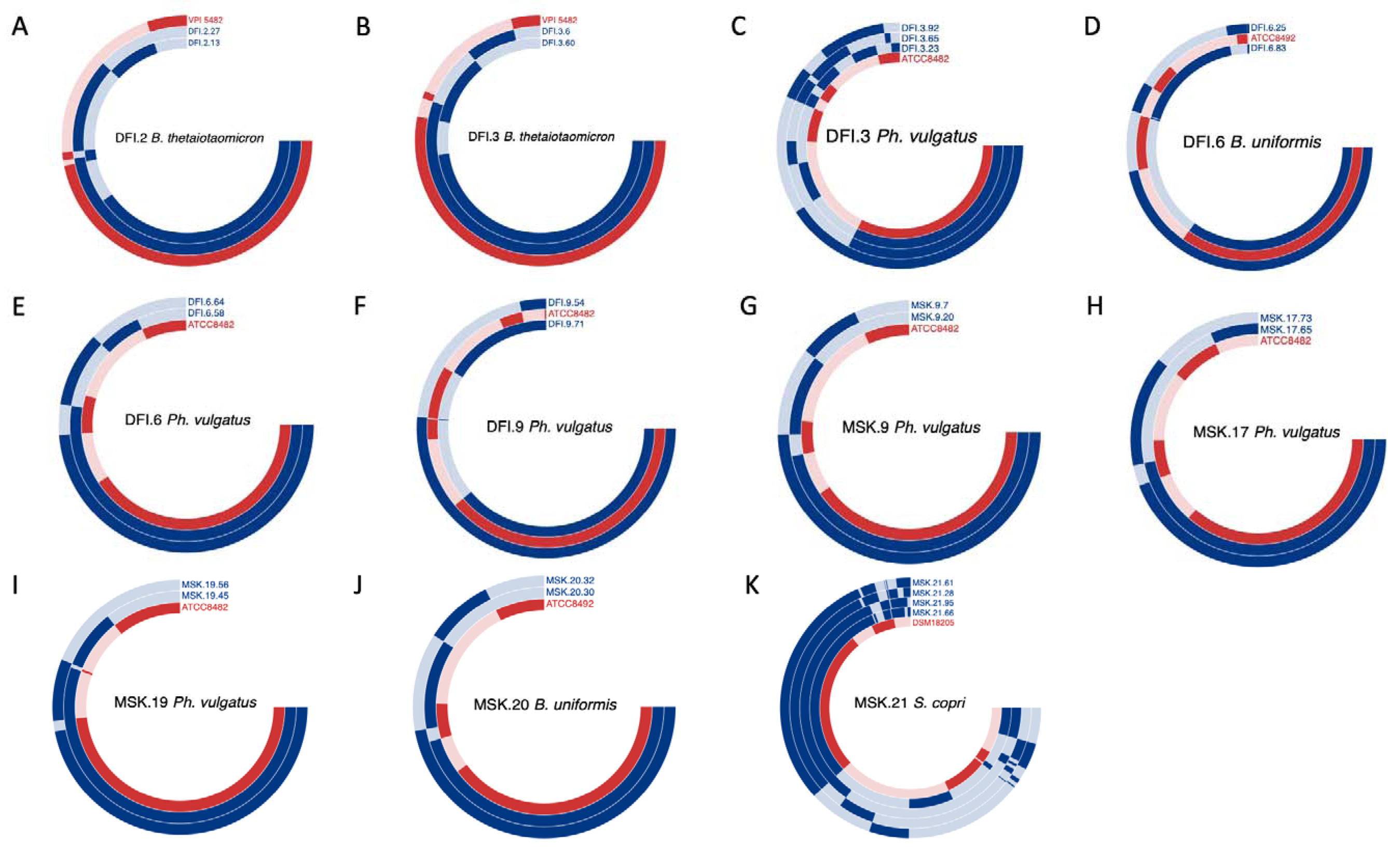
Pan-genome comparison of strains of the same species from the same donor. Pangenome comparison circular plot among Bacteroides thetaiotaomicron strains from donor DFI.2 (A), B. thetaiotaomicron strains from donor DFI.3 (B), Phocaeicola vulgatus strains from donor DFI.3 (C), Bacteroides uniformis strains from donor DFI.6 (D), Ph. vulgatus strains from donor DFI.6 (E), Ph. vulgatus strains from donor DFI.9 (F), Ph. vulgatus strains from donor MSK.9 (G), Ph. vulgatus strains from donor MSK.17 (H), Ph. vulgatus strains from donor MSK.19 (I), B. uniformis strains from donor MSK.20 (J), and Segatella copri strains from donor MSK.21 (K). Corresponding type strains are also included in red tracks. Light shading indicates absence of that gene cluster.

**Figure S4.**
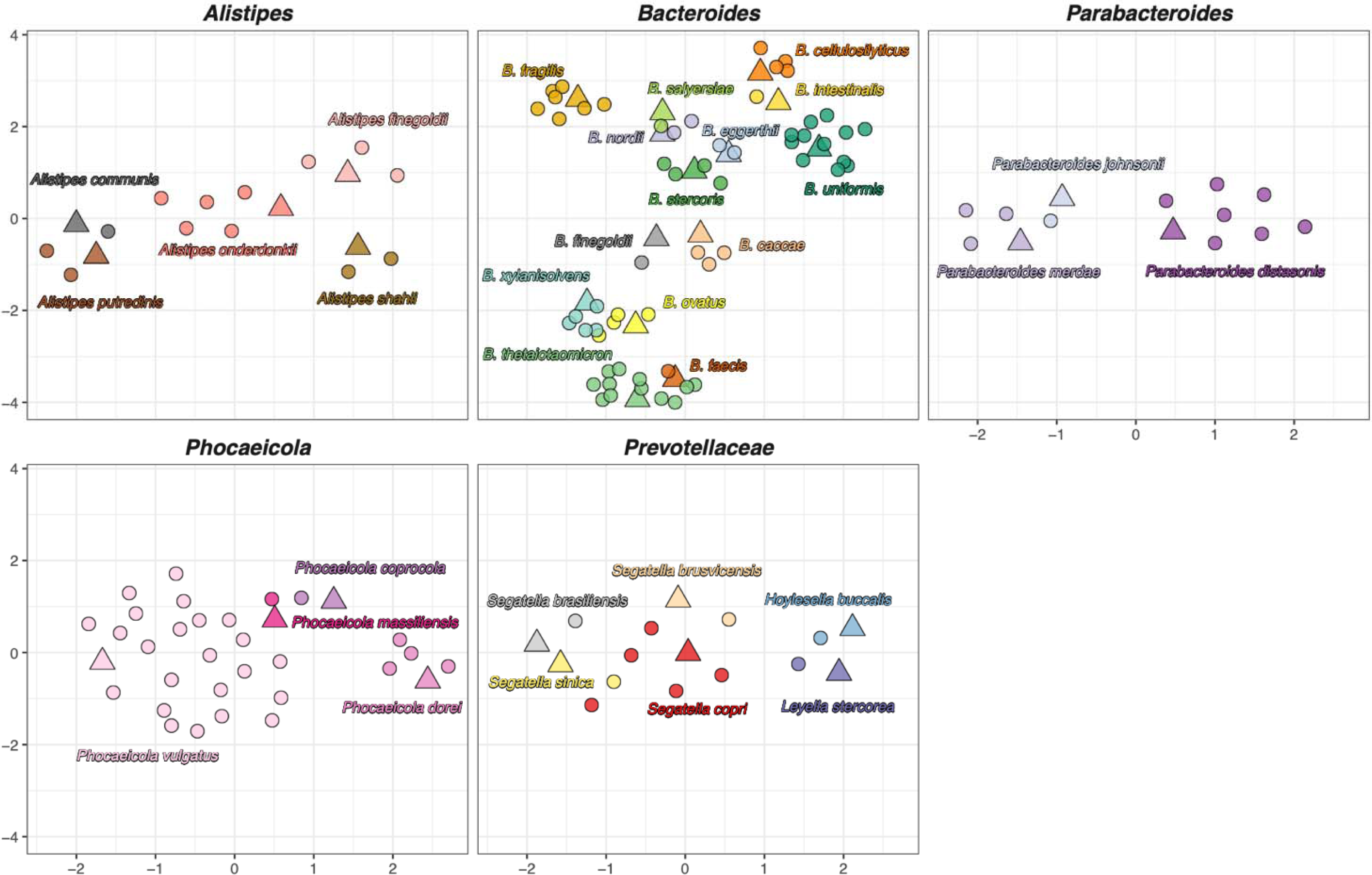
UMAP analysis of the genomic diversity of Bacteroidales isolates within the genera or family. Whole-genome gene cluster UMAP based on presence or absence of gene clusters in Bacteroidales genera and families. Triangle indicates type strains, whereas circles indicate human isolates included in our collection.

**Figure S5.**
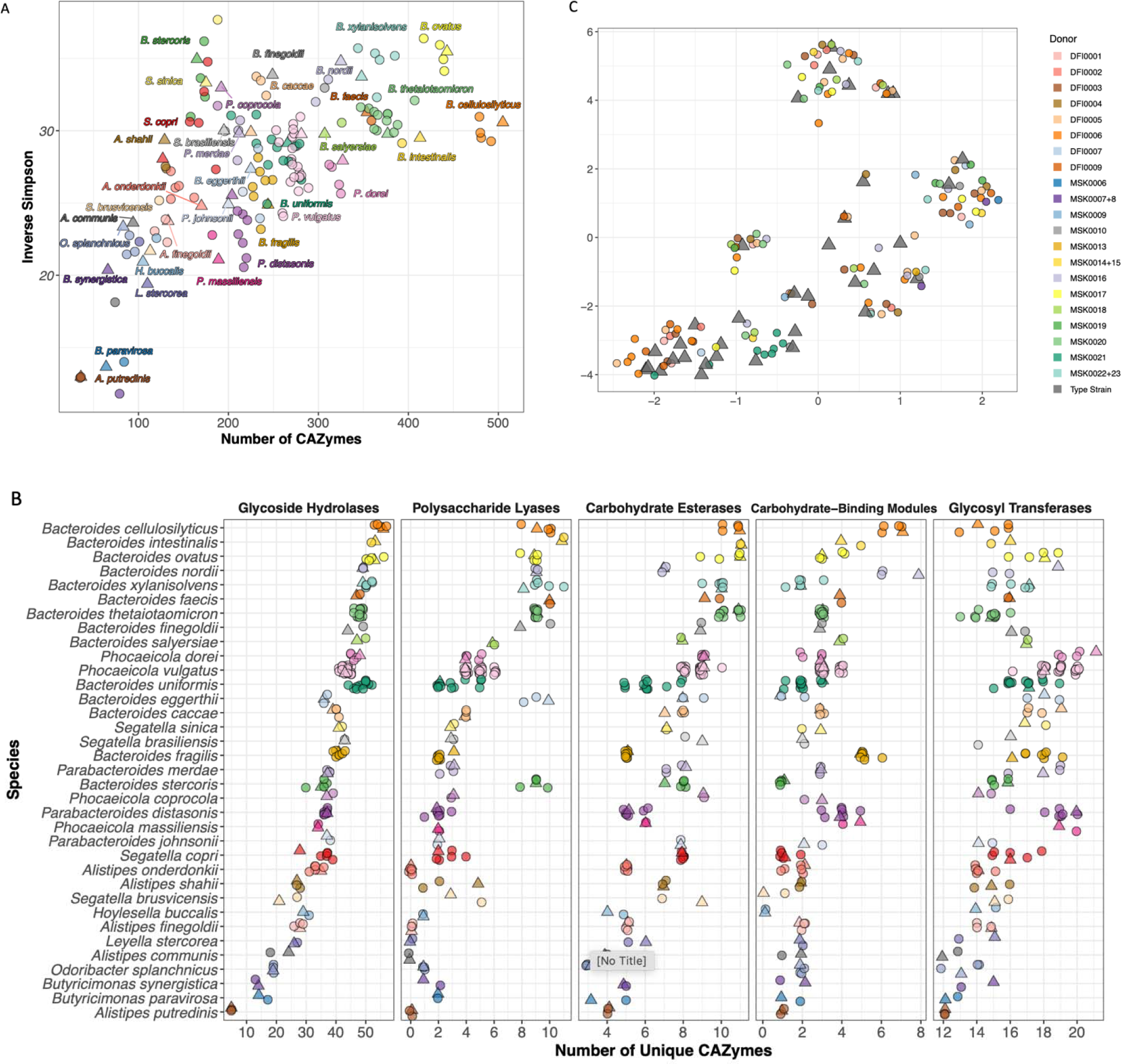
Additional CAZyme and PUL analyses of Bacteroidales isolates. A. Number of CAZymes in Bacteroidales isolates and their Inverse Simpson Indices. B. Numbers of unique CAZyme families in each Bacteroidales isolate, arranged by species from most (top) to least (bottom). CAZyme families are categorized into five major classes according to CAZy database, each panel representing one class. C. UMAP plot based on the presence or absence of CAZyme families in all Bacteroidales isolates and type strains (as in Figure 4B). Color indicates donor identity.

**Figure S6.**
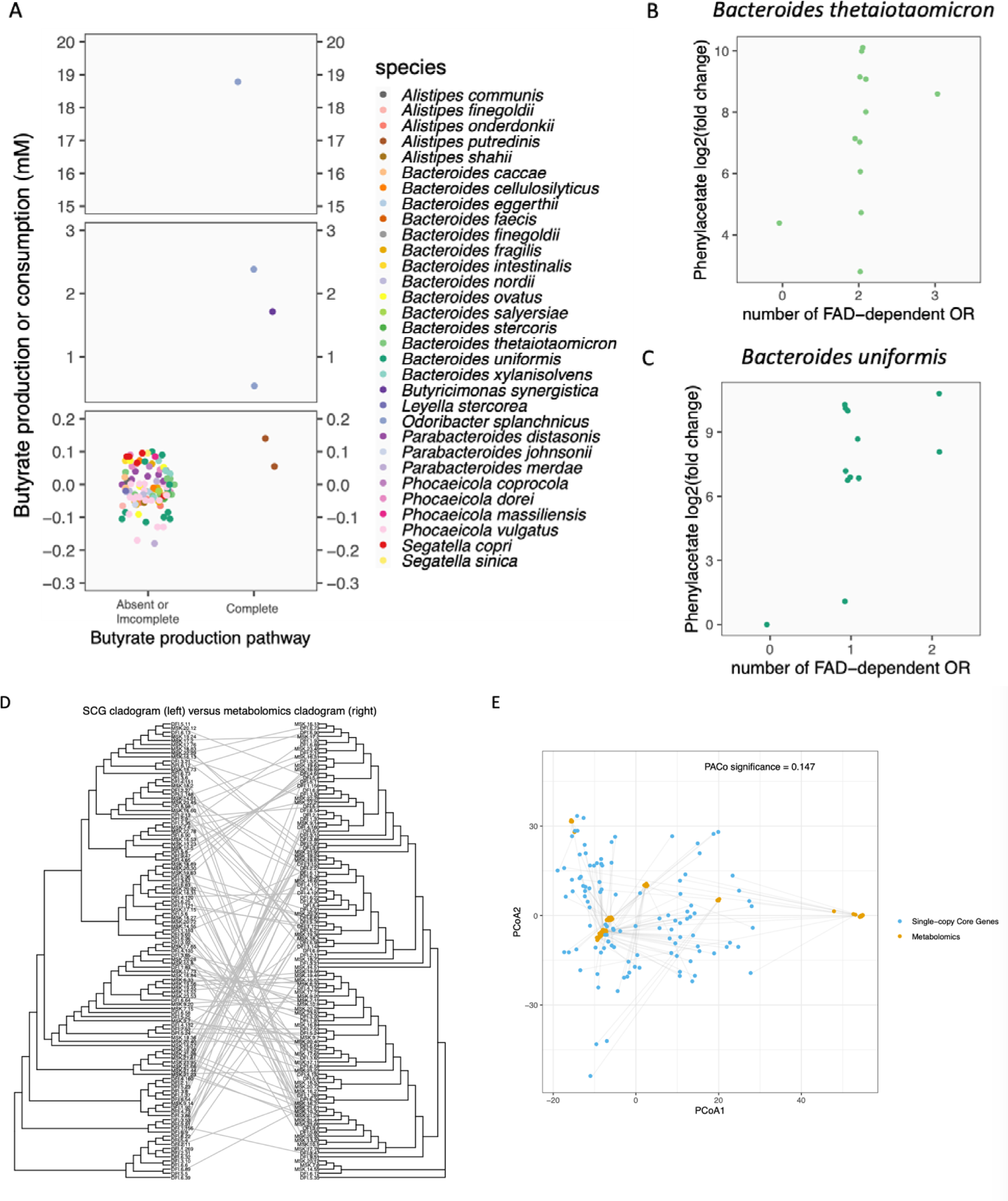
Inter- and intra-specific diversity of metabolome among Bacteroidales isolates. A. Butyrate concentration changes after culturing in BHIS among Bacteroidales isolates with complete or incomplete/absent butyrate production pathways in their genomes. Color indicates species identity. B. Relative concentration fold change of phenylacetate after culturing in BHIS among Bacteroides thetaiotaomicron isolates compared to numbers of FAD-dependent oxidoreductase (OR) in their genomes. C. Relative concentration fold change of phenylacetate after culturing in BHIS among Bacteroides uniformis isolates compared to numbers of FAD-dependent oxidoreductase (OR) in their genomes. D. Comparison between single-copy core gene dendrogram and hierarchical clustering dendrogram of metabolomic profiles of Bacteroidales isolates. Lines connect the same isolate on two dendrograms. E. Procrustes superimposition of PCoA plots of single-copy core genes (blue) and metabolomic profiles (orange). Lines connect the same isolate on two PCoA plots.

**Figure S7.**
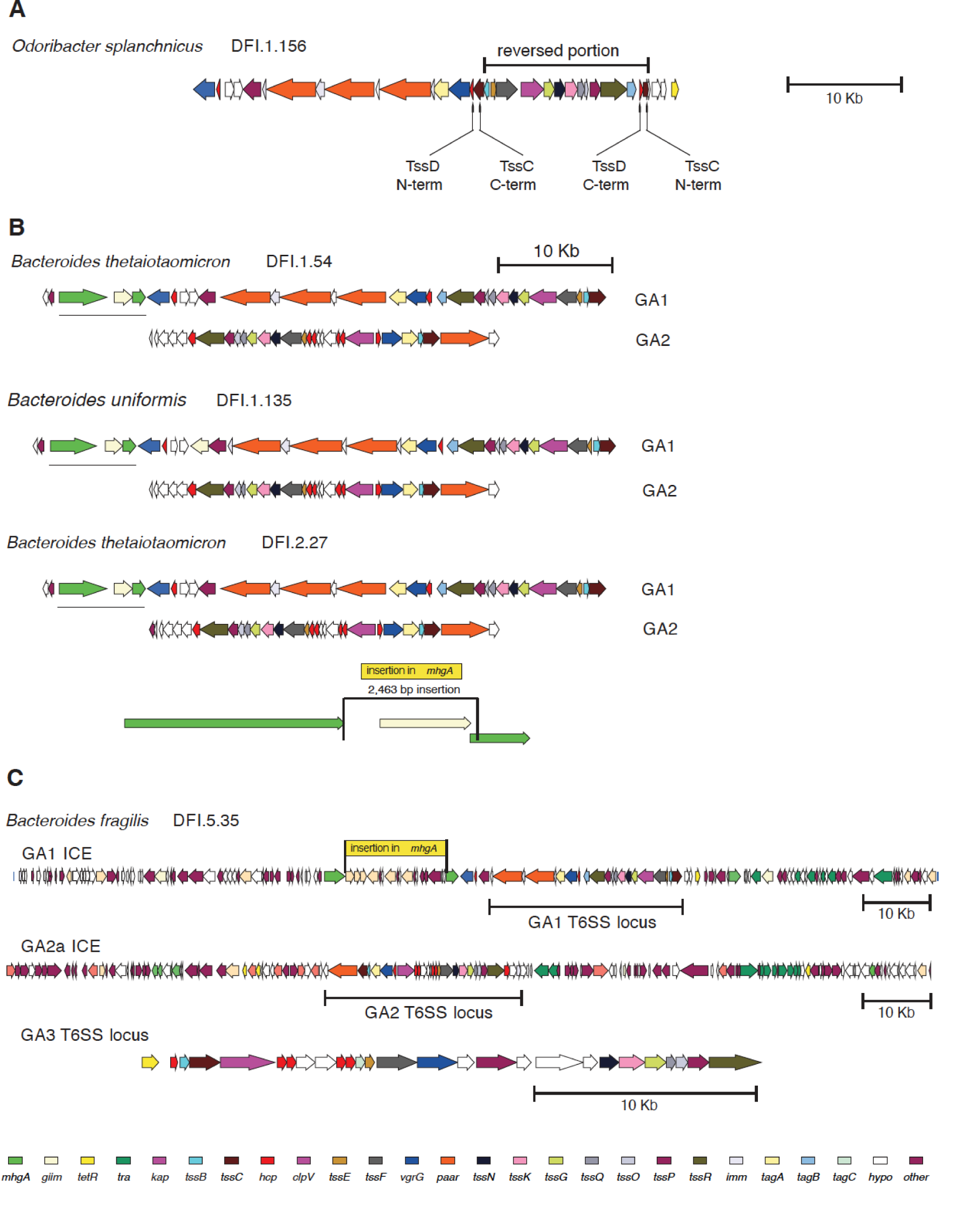
Genetic features of select Type VI secretion system loci among Bacteroidales isolates. A. A dysfunctional GA1 T6SS locus from Odoribacter splanchnicus where 14 of the tss genes are inverted resulting in partial tssD and tssC genes at the inversion termini. B. Open reading frame maps of the GA1 and flanking regions showing the mhgA gene (green) interrupted by a 2463 bp insertion encoding a transposase-like gene in strains also containing a GA2 locus. C. The T6SS genetic regions of a B. fragilis strains containing all three T6SS genetic architectures (GA1-GA3). The mghA gene near the GA1 locus is also interrupted but by a much larger insertion.

