## Supplementary material for "Comprehensive analyses of a large human gut Bacteroidales culture collection reveal species and strain level diversity and evolution": Methods

### Methods for

##### **Authors and Affiliations**

**Duchossois Family Institute, University of Chicago, 900 E. 57th St, Chicago, IL, 60637, USA**

Zhenrun J. Zhang, Huaiying Lin, Michael J. Coyne, Cody G. Cole, Nicholas Dylla, Rita C. Smith, Emily Waligurski, Ramanujam Ramaswamy, Victoria Burgo, Che Woodson, Jessica C. Little, David Moran, Amber Rose, Mary McMillin, Emma McSpadden, Anitha Sundararajan, Ashley M. Sidebottom, Eric G. Pamer, Laurie E. Comstock

**Department of Medicine, Section of Infectious Diseases & Global Health, University of Chicago Medicine, 5841 South Maryland Ave, Chicago, IL, 60637, USA**

Eric G. Pamer

**Department of Microbiology, Biological Sciences Division, University of Chicago, 5841 South Maryland Ave, Chicago, IL, 60637, USA**

Zhenrun J. Zhang, Michael J. Coyne, Cody G. Cole, Emily Waligurski, Eric G. Pamer, Laurie E. Comstock

### Methods

#### Fecal sample donors

Twenty-one healthy human donors were enrolled in a prospective fecal collection protocol. Donor age and sex were not reported. The prospective fecal collection protocol was approved by the institutional review boards at Memorial Sloan Kettering Cancer Center and at The University of Chicago. All donors provided written and informed consent for IRB-approved biospecimen collection and analysis (protocols 06-107; IRB20-1384). The study was conducted in accordance with the Declaration of Helsinki.

#### Isolation and growth of Bacteroidales isolates

Isolation and growth of Bacteroidales isolates are similar to previously described.^1^ Isolation and growth of commensal bacteria was performed under anaerobic conditions with 3% H_2_, 5% CO_2_ and 92% N_2_ in an anaerobic chamber (Coy Labs). Fresh donor fecal samples were transferred into anaerobic conditions within 1 h of collection. Fecal samples were resuspended in pre-reduced PBS and plated on Columbia agar with 5% sheep blood (BBL, BD) or brain-heart infusion (BHI – Difco, BD) agar in three serial 10-fold dilutions and incubated at 37°C for 48 – 96 h. Isolated colonies were re-streaked for purity onto Columbia blood agar and frozen in pre-reduced 10% glycerol in PBS. For broth cultures, isolates were grown in pre-reduced BHI supplemented with 5g/L yeast extract (Difco, BD) and 0.1% L-cysteine (Sigma) (BHIS).

#### Whole-genome shotgun sequencing and assembly

Bacterial DNA was extracted using the QIAamp PowerFecal Pro DNA kit (Qiagen). Prior to extraction, samples were subjected to mechanical disruption using a bead beating method. Briefly, samples were suspended in a bead tube (Qiagen) along with lysis buffer CD1 and loaded on a bead mill homogenizer (Fisherbrand). Samples were then centrifuged, and supernatant was resuspended in CD2, a reagent that effectively removes inhibitors by precipitating non-DNA organic and inorganic materials including polysaccharides, cell debris and proteins. DNA was then purified routinely using a spin column filter membrane and quantified using Qubit (Life Technologies). Libraries were prepared using 200 ng of genomic DNA using the QIAseq FX DNA library kit (Qiagen). Briefly, DNA was fragmented enzymatically using a nuclease into shorter fragments and desired insert size was achieved by adjusting fragmentation conditions. Fragmented DNA was end repaired to generate blunt end fragments using a T4 DNA polymerase, and ‘A’s’ were added to the 3’ends to stage inserts for ligation. During ligation step (blunt end AT ligation), Illumina compatible Unique Dual Index (UDI) adapters were added to the inserts and prepared library was PCR amplified. Amplified libraries were recovered using magnetic beads, and QC was performed using Tapestation 4200 (Agilent Technologies). Libraries were sequenced on an Illumina MiSeq or NextSeq 500 platforms to generate 2x250bp or 2x150bp reads, respectively, targeted for 1 to 3 million pair-end reads per sample. Adapters were trimmed with Trimmomatic^2^ using following parameters: the leading and trailing 3 bp of the sequences were trimmed off, quality was controlled by a sliding window of 4, with an average quality of 15. Moreover, any read that was less than 50 bp long after trimming and quality control were discarded. The remaining high-quality reads were assembled into contigs using SPAdes (v.3.14.0).^3^

#### 16S rRNA sequencing and analysis

The V4-V5 region of the 16S rRNA gene was amplified using universal bacterial primers – 563F (5’-nnnnnnnn-NNNNNNNNNNNN-AYTGGGYDTAAA-GNG-3’) and 926R (5’-nnnnnnnn-NNNNNNNNNNNN-CCGTCAATTYHT-TTRAGT-3’), where ‘N’ represents the barcodes, ‘n’ are additional nucleotides added to offset primer sequencing. The approximately ~412 bp amplicons were then purified using a spin column-based method (Minelute, Qiagen), quantified, and pooled at equimolar concentrations. Illumina sequencing-compatible Combinatorial Dual Index (CDI) adapters were ligated onto the pools using the QIAseq 1-step amplicon library kit (Qiagen). Library QC was performed using Qubit and Tapestation and sequenced on Illumina MiSeq platform to generate 2x250bp paired-end reads. For 16S rRNA amplicon sequence analysis, we used DADA2 (v1.18.0)^4^ as our default pipeline for processing MiSeq 16S rRNA reads with minor modifications in R (v4.0.3). Specifically, reads were first trimmed at 210 bp for forward reads and 150 for reverse reads to remove low quality nucleotides. Chimeras were detected and removed using the default consensus method in the DADA2 pipeline. Then, ASVs with length between 300 bp and 360 bp were kept and deemed as high quality ASVs. Taxonomy of the resultant ASVs were assigned to the genus level using the RDP classifier (v2.13)^5^ with a minimum bootstrap confidence score of 80. Results were collapsed to the Genus level and plotted using R and the ggplot2 package.

#### Nanopore sequencing and assembly

Samples for Nanopore and Illumina hybrid assemblies were extracted using the NEB Monarch Genomic DNA Purification Kit or the QIAamp PowerFecal Pro DNA kit (Qiagen). DNA was quality checked using a genomic Tapestation 4200. Nanopore libraries were prepared using the Ligation Sequencing Kit (SQK-LSK109), the Native Barcoding Expansions 1-12 (EXP-NBD104) and 13-24 (EXP-NBD114), and the NEBNext Companion Module for Oxford Nanopore Technologies (E7180S). The shearing steps and first ethanol wash were eliminated to ensure high concentrations of long fragments. Using R9.4.1 flow cells, libraries were run on a MinION for 72 hours at ≈180mV. Hybrid assemblies were completed using Unicycler (v0.4.8)^6^ with default parameters, using the non-filtered Nanopore long reads with and without prior trimming of the Illumina short reads as input. The short reads were trimmed by Trim Galore (v.0.4.5) with following parameters: the adapter and the leading 6 bp of forward and reverse short reads were trimmed off, the quality was controlled at 30, while minimum length was at 75 bp.

#### De-replication of Bacteroidales isolates

1335 genomes deposited to RefSeq^7^ under BioProject IDs PRJNA737800 and PRJNA792599 were retrieved, and 408 Bacteroidales genomes from donors comprising the DFI and MSK cohorts were retained (Dataset 1, Bacteroidales collection tab). Isolates arising from sampling of the same subject (MSK.7 & MSK.8, MSK.14 & MSK.15, and MSK.22 & MSK.23) were combined into single communities. Community MSK.5 was eliminated due to the health status of the donor at the time of sample collection. Additionally, *Bacteroides eggerthii* strains DFI.5.6A and DFI.5.6B were removed as direct duplicates of *Bacteroides eggerthii* DFI.5.6, and *Bacteroides thetaiotaomicron* strains DFI.3.60 and DFI.3.60A were removed as direct duplicates of *Bacteroides thetaiotaomicron* DFI.3.60B. Speciation of the remaining isolates was confirmed and/or assigned using a variety of methods, including 16S sequence analysis, the GTDB project,^8^ comparison of single-copy core genome genes (see Dataset 1, tab 25 core genes and Figure S3) using Anvi'o,^9^ and comparison to type strain genomes retrieved from RefSeq. Assistance with speciation of the *Segatella* species isolates was generously provided by Prof. Dr. Till Strowig and his group at the Helmholtz Centre for Infection Research, Braunschweig, Germany (personal communication) and reference.^10^ Isolates arising from the same clonal lineage within a donor community were identified by first comparing genome sequences using the dnadiff.pl script from MUMmer v. 4.0.0^11^ and utilizing threshold cutoffs for the AvgIdentity (> 99.9%), TotalGSNPs (< 1000), and TotalGIndels (< 75) outputs of each comparison. Further analysis was performed using FastANI.^12^ These analyses informed our final set of 408 Bacteroidales genomes comprising 127 unique clonal lineages from 21 communities (see Datset 1, de-duplication tab).

#### Phylogenetic tree and taxonomic classification of Bacteroidales isolates

For phylogenetic comparisons, type strains of the species included in the analyses that were isolated from humans were retrieved from RefSeq database. The phylogenetic tree was constructed using single-copy core genes (SCGs) identified during the Anvi’o pangenome bioinformatic workflow^13^ through Hidden Markov Models (HMM) of a list of 71 bacterial SCGs. from this list, 25 genes were indeed present in a single copy and found in all the genomes including the *Escherichia coli* K12 MG1655 outgroup used for rooting the tree (Dataset 1, tab 1). The protein sequences encoded for by these genes were extracted and concatenated using the `anvi-get-sequences-for-hmm-hits`. A Newick file was generated using the `anvi-gen-phylogenomic-tree` function, which utilizes FastTree 2.1.11.^14^ Taxonomic classification of the isolates is based on their closest type strain, corroborated by two other methods: (a) full/partial length 16S rRNA gene from each isolate are queried using BLASTn (v2.10.1+)^15^ against NCBI’s 16S ribosomal RNA sequences database (dated Nov 2020), and top five hits for each query are manually curated to determine an isolate’s identity, with identity and coverage cutoff at 95%; (b) GTDB-Tk (v1.5.1).^16^

#### Core genome analysis

Anvi’o^9^ was used for core genome analysis. Gene calls were imported from the GenBank files generated after custom Prokka annotation using `anvi-script-process-genbank`. Processed GenBank files were then analyzed using the Anvi’o pangenome bioinformatic workflow with the function `anvi-run-workflow`. In the workflow, NCBI’s BLAST was selected for quantification of amino acid sequence similarity. To reduce noise from weak matches between amino acid sequences, the default 0.5 minbit heuristic parameter was used. Protein clusters were identified from remaining amino acid sequence similarities using the Markov clustering algorithm (MCL) with an inflation parameter of 2. Clustering results were exported using the `anvi-summarize` function. Protein clusters were matched for each genome at the family, genus, species, and strain level taxonomy and plotted using R.

#### Strain pangenome comparison

Pangenomes of strains of the same species from the same donor were generated using an Anvi’o workflow similar to the core genome analysis (above), but with a Markov clustering algorithm (MCL) inflation parameter of 10. Similarity between strains was determined as the number of shared unique protein clusters divided by the total unique protein clusters between the strains being compared.

#### UMAP analysis of whole-genome genetic repertoire

The global repertoire of all protein clusters was counted as being either present or absent in each of the Bacteroidales isolates. A uniform manifold approximation and projection (UMAP) analysis was used to group isolates using a Manhattan distance metric.^17^ Similar analyses were performed on select genera and families.

#### Genomic annotation

For individual isolates, the genome assemblies were annotated using Prokka (v. 1.12)^18^ modified to utilize additional HMM models, including Pfam35 (November 2021)^19^, TIGRFAMs (v. 15),^20^ COGs (2020 version),^21^ and the previously described T6SS and Toxin & Immunity HMM model sets.^22^

#### Analyses of CAZymes and PULs

For genomic annotation of CAZymes, hidden Markov models of CAZyme families and subfamilies from dbCAN database^23^ were used as templates for mining among genomes using hmmscan in HMMER3^24^ with a seq_evalue cutoff of 1e-18. A uniform manifold approximation and projection UMAP analysis^17^ was used to group isolates according to the presence or absence of CAZy families and subfamilies. PULpy^25^ was used with default settings for mining PULs in genomes. SusCD-only PULs were excluded unless otherwise noted.

#### Analysis of T6SSs and antibacterial genes

Detection of T6SS loci utilized the concatemer sequences for each T6SS genetic architecture delineated in reference^26^ as queries for a BLASTn analysis against our database of 408 Bacteroidales genomes. Best hit analysis of the BLASTn returns is tabulated in the T6SS-antibacterial genes tab of Dataset 1. Protein sequences of bacteroidetocin A (Bd-A) and bacteroidetocin B (Bd-B)^27^, along with that of the *Bacteroides fragilis* Ubb protein (BfUbb)^28^ were used in a tblastn search against the same database, and the results were compiled. Protein sequences of BSAP-1,^29^ BSAP-2,^30^ BSAP-3,^31^ BSAP-4,^32^ BcpT,^33^ and the fragipain-activated bacteriocin Fab1^34^ were used in a blastp analysis and used a evalue cutoff of 1e-50. Additionally, a scan for the presence of a globally distributed cryptic plasmid of the human gut, pBI143,^35^ was performed using blastn; these data and the thresholds employed are included in Dataset 1.

#### Inter-species DNA transfer analyses

DNA regions conserved between isolates from the same communities were detected using the blastn program from the BLAST v 2.13.0+ suite,^15^ with settings -perc_identity = 99.99, -evalue = 1e-50, and with both -dust and soft masking settings disabled. The BLASTn output was further parsed to require a minimum alignment length of 5000 bp. The segments identified by BLASTn were retrieved and those from the same community were clustered using CD-HIT v 4.8.1.^36^ The CD-HIT output was parsed to detect the cluster representatives, and these were used as representative samples of the conserved regions. Annotation and CDS region coordinates for clusters detected in more than one species within the same community were retrieved, and the per-community lists were manually curated (see Dataset 3).

#### Intra-species DNA difference analyses

Novel regions of DNA within clonal isolates (see Dataset 1, de-duplication tab) were detected using the standalone version of PanSeq v 3.2.1^37^ run locally under Red Hat Enterprise Linux v. 9 in its “novel” run mode, with settings minimumNovelRegionSize and novelRegionFinderMode set to 5000 and no_duplicates, respectively. Also utilized in these analyses were MUMmer v. 4.0.0,^11^ BLAST v 2.13.0+,^15^ Muscle v. 5.1,^38^ and CD-HIT v 4.8.1.^36^ The genomic sequence of each clonal isolate was compared in a reciprocal manner. The novel regions found were then clustered using CD-HIT. The annotation and CDS region data were retrieved for each novel region, and manually curated to further reduce redundancy and detect regions that were likely in close proximity (see Dataset 4).

#### Bile acid modification pathway analyses

Several protein queries for each of seven bile acid modification pathway enzymes (3β-HSDH-I, 3β-HSDH-II, 7α-HSDH, 5AR, 5BR, Bs_hydrolase-I, Bs_hydrolase-II, Bs_hydrolase-III)^39,40^ were utilized in a BLASTp analysis (evalue threshold = 1e-50) using the protein sequences compiled from our collection of 408 Bacteroidales genome (Dataset 1, Bacteroidales collection tab). The BLASTp output was parsed to detect the best hit for each query group for each genome and tabulated.

#### Metabolomic profiling

Bacteroidales isolates were streaked from stocks in PBS with 10% glycerol onto Colombia blood agar plates (BD, B21263X) and incubated at 37°C for 24-48 hours anaerobically, until distinct colonies formed. One to three colonies were suspended in 200 µL Bacto Brain Heart Infusion Media (BD, 237500, lot 0307917) as inoculum, which was further diluted 10-fold into 300 µL of Bacto Brain Heart Infusion Media in 96 well non-TC plates (Fisher Sci, 7000108) with duplicates. The plates were loaded into a Logphase 600 plate reader (BioTek, BTLP600) and incubated at 37°C with double orbital shaking at 500 rpm. Incubation period ranged from 16-72 hours, depending on the growth rates of the strains being profiled. Strains were grown until OD readings indicated late log or early stationary phase had been achieved. Duplicate wells were pooled into a single well of a 48-well non-TC plate (Fisher Sci, 877252) and centrifuged at 4300 RCF for 15 minutes. Avoiding the pellet, 50-100 µL of the supernatants were transferred into a new 48-well non-TC plate and sealed using a plate film. BHI media was included as baseline control. These supernatants were frozen at -80°C until further analysis.

Supernatants were extracted with extraction solvent (4 volumes of 100% methanol spiked with internal standards and stored at -80°C) in a microcentrifuge tube. Tubes were then centrifuged at -10°C, 20,000 x g for 15 min and supernatant was used for subsequent metabolomic analysis. Metabolites were derivatized as described by Haak *et al.* with the following modifications^41^. The metabolite extract (100 μL) was added to 100 μL of 100 mM borate buffer (pH 10) (Thermo Fisher, 28341), 400 μL of 100 mM pentafluorobenzyl bromide (Millipore Sigma; 90257) in acetonitrile (Fisher; A955-4), and 400 μL of *n*-hexane (Acros Organics; 160780010) in a capped mass spec autosampler vial (Microliter; 09-1200). Samples were heated in a thermomixer C (Eppendorf) to 65°C for 1 hour while shaking at 1300 rpm. After cooling to room temperature, samples were centrifuged at 4°C, 2000 x g for 5 min, allowing phase separation. The hexanes phase (100 μL) (top layer) was transferred to an autosampler vial containing a glass insert and the vial was sealed. Another 100 μL of the hexanes phase was diluted with 900 μL of *n*-hexane in an autosampler vial. Concentrated and dilute samples were analyzed using a GC-MS (Agilent 7890A GC system, Agilent 5975C MS detector) operating in negative chemical ionization mode, using a HP-5MSUI column (30 m x 0.25 mm, 0.25 μm; Agilent Technologies 19091S-433UI), methane as the reagent gas (99.999% pure), and 1 μL split injection (1:10 split ratio). Oven ramp parameters were as follows: 1 min hold at 60°C, 25°C per min up to 300°C with a 2.5 min hold at 300°C. Inlet temperature was 280°C and transfer line was 310°C. For quantitative metabolomics of SCFAs, a 10-point calibration curve was prepared with acetate (100 mM), propionate (25 mM), butyrate (12.5 mM), and succinate (50 mM), with 9 subsequent 2x serial dilutions. Data analysis was performed using MassHunter Quantitative Analysis software (version B.10, Agilent Technologies) and confirmed by comparison to authentic standards. Normalized peak areas were calculated by dividing raw peak areas of targeted analytes by averaged raw peak areas of internal standards, which were then converted to quantitative values in mM concentration. For semi-quantitative metabolomics, values were calculated from the raw peak area of the compound normalized to the median raw peak area of two baseline controls (BHI media).

The presence of butyrate production pathways and Gut Metabolic Modules (GMMs) among isolates were mapped using Omixer-RPM in GOmixer.^42,43^ Counts of FAD-dependent oxidoreductase in Bacteroidales genomes were tallied based on Prokka annotation. Hierarchical clustering of metabolomic profiles of isolates was performed with pheatmap package in R.

For Procrustes cophylogeny test between phylogeny and metabolomy, pairwise distance matrix of metabolomy of isolates were calculated based on their semi-quantitative metabolomics. Pairwise distance matrix of phylogeny of these isolates were calculated based on their SCG phylogenetic tree. Procrustes cophylogeny test between phylogeny and metabolonomy was performed using PACo (v0.4.2),^44^ using distance matrix of phylogeny as “host” and distance matrix of metabolomy as “parasite”. PACo test was performed with row and column swapping (method = ‘r00’) and 1000 permutations (nperm = 1000).
